# Atypical B cells consist of subsets with distinct effector functions

**DOI:** 10.1101/2022.09.28.509955

**Authors:** Raphael A. Reyes, Gayani Batugedara, Paramita Dutta, Ashley B. Reers, Rolando Garza, Isaac Ssewanyana, Prasanna Jagannathan, Margaret E. Feeney, Bryan Greenhouse, Sebastiaan Bol, Ferhat Ay, Evelien M. Bunnik

## Abstract

Atypical B cells are a population of activated B cells that are commonly enriched in individuals with chronic immune activation, but are also part of a normal immune response to infection or vaccination. Prior studies to determine the function of these cells have yielded conflicting results, possibly due to functional heterogeneity among this B cell population. To better define the role(s) of atypical B cells in the host adaptive immune response, we performed single-cell sequencing of transcriptomes, cell surface markers, and B cell receptors in individuals with chronic *Plasmodium falciparum* exposure, a condition known to lead to accumulation of circulating atypical B cells. Our studies identified three previously uncharacterized populations of atypical B cells with distinct transcriptional and functional profiles, that separate into two differentiation pathways. We identified a set of cell surface markers to distinguish these atypical B cell subsets and confirmed their presence in malaria-experienced children and adults using flow cytometry. *Plasmodium falciparum*-specific cells were present in equal proportions within each of these atypical B cell populations, indicating that all three subsets develop in response to antigen stimulation. However, we observed marked differences among the three subsets in their ability to produce IgG upon T-cell-dependent activation. Collectively, our findings help explain the conflicting observations in prior studies on the functions of atypical B cells and provide a better understanding of their role in the adaptive immune response in chronic inflammatory conditions.

**One sentence summary:** Atypical B cells consist of three subsets that may play distinct roles in the host adaptive immune response.

## INTRODUCTION

Humoral immunity to infection relies on the development of memory B cells that circulate through the blood and secondary lymphoid organs to patrol for antigen, as well as long-lived plasma cells that home to the bone marrow and function as a source of antibodies that inhibit pathogen replication and survival. In humans, memory B cells are defined based on the expression of the surface proteins CD21 and CD27. Upon vaccination or acute infection, other antigen-experienced B cell populations transiently increase in abundance in the circulation, including CD21^-^CD27^+^ activated B cells and CD21^-^CD27^-^ atypical B cells (also called DN2 cells) (*1–3*). These activated and atypical B cell populations are thought to be part of a normal immune response and peak in abundance within 2 to 4 weeks after antigen exposure, followed by a gradual return to baseline over the course of several months (*1, 4*). However, under conditions of chronic or repetitive immune activation, as seen during HIV or *Plasmodium* infection, and in autoimmune diseases, such as systemic lupus erythematosus (SLE), atypical B cells accumulate and become a more prominent presence in the circulation (*5–8*). In healthy adults, atypical B cells make up approximately 2 - 4% of all circulating B cells, but this can increase to 20 – 40% in individuals with chronic inflammatory conditions (*5, 6, 9*). Whether these atypical B cells contribute to control of infections or negatively affect the host immune response remains incompletely understood. A better understanding of the function of atypical B cells will provide insight into the immune response to chronic infection and may present ways to overcome immune dysfunction or to harness these cells by vaccination.

Studies regarding the functionality of atypical B cells in the immune response have highlighted various differences between atypical B cells and memory B cells. Specifically, atypical B cells display reduced expression of B cell receptor signaling pathway genes and costimulatory molecules, as well as upregulation of inhibitory receptors (*10, 11*). In addition, atypical B cells were less responsive to B cell receptor engagement by soluble antigen and showed reduced differentiation into antibody-secreting cells under conditions that efficiently induced differentiation of memory B cells into antibody-secreting cells (*5, 8-10, 12*). These results initially led to the conclusion that atypical B cells are exhausted and non-responsive to stimulation. However, it was recently reported that atypical B cells respond robustly to high-affinity membrane-associated antigen and showed a markedly reduced response to low-affinity antigens, a selective mechanism which may serve to limit activation of atypical B cells by low-affinity antigens, such as auto-antigens (*11, 13*). Despite these new findings, the contribution of atypical B cells to the host immune response remains subject of debate and various functions have been proposed for these cells. These include (pre-)antibody secreting cells, based on the presence of secretory Ig transcripts and the overlap in B cell receptor sequences between atypical B cells and plasma cells, and immune-modulatory cells, based on the upregulation of surface proteins that promote interactions between B cells and T cells (*9, 11, 14-18*). The multiple and conflicting effector functions previously assigned to atypical B cells are suggestive of functional heterogeneity among this population.

Here, we used chronic *P. falciparum* exposure as a model to define the heterogeneity of the atypical B cell compartment. Over the course of many years of repetitive *P. falciparum* infections, people living in malaria-endemic regions develop an immune response that protects against disease. As a result, most cases of malaria occur in children under the age of 10, while adults with life-long exposure have asymptomatic infections. To study atypical B cells induced by malaria, we performed single-cell transcriptomic analysis with cell surface marker and B cell receptor profiling on longitudinal samples obtained from two children following malaria. Based on these sequencing data, we defined three subsets of atypical B cells with distinct transcriptional profiles that are likely to play different roles in the immune response. Using flow cytometry, we then analyzed the longevity and *Plasmodium* antigen-specificity among these subsets in samples from malaria-experienced children and adults.

## RESULTS

### Single-cell transcriptomics identifies B cell populations

The atypical B cell population is strongly expanded in the first few weeks after a malaria episode and slowly contracts to baseline levels over the course of several months (*4*). To study the identity of atypical B cells induced as a result of the hyperinflammatory immune response during malaria and following return to baseline conditions, we selected two children (5 and 7 years old) for whom peripheral blood mononuclear cell (PBMC) samples were collected three weeks after malaria and six months later (**Figure 1A**, **Table S1**). Both children were free of symptomatic malaria between the two time points, but had detectable *P. falciparum* parasitemia at both the three- and six-month time points (**Figure S1**). Asymptomatic *P. falciparum* infections result in very low immune activation (*19*), and it can therefore be expected that the atypical B cell population is only minimally affected by these infections, especially in comparison to the strong inflammatory response during malaria.

**Figure 1:**
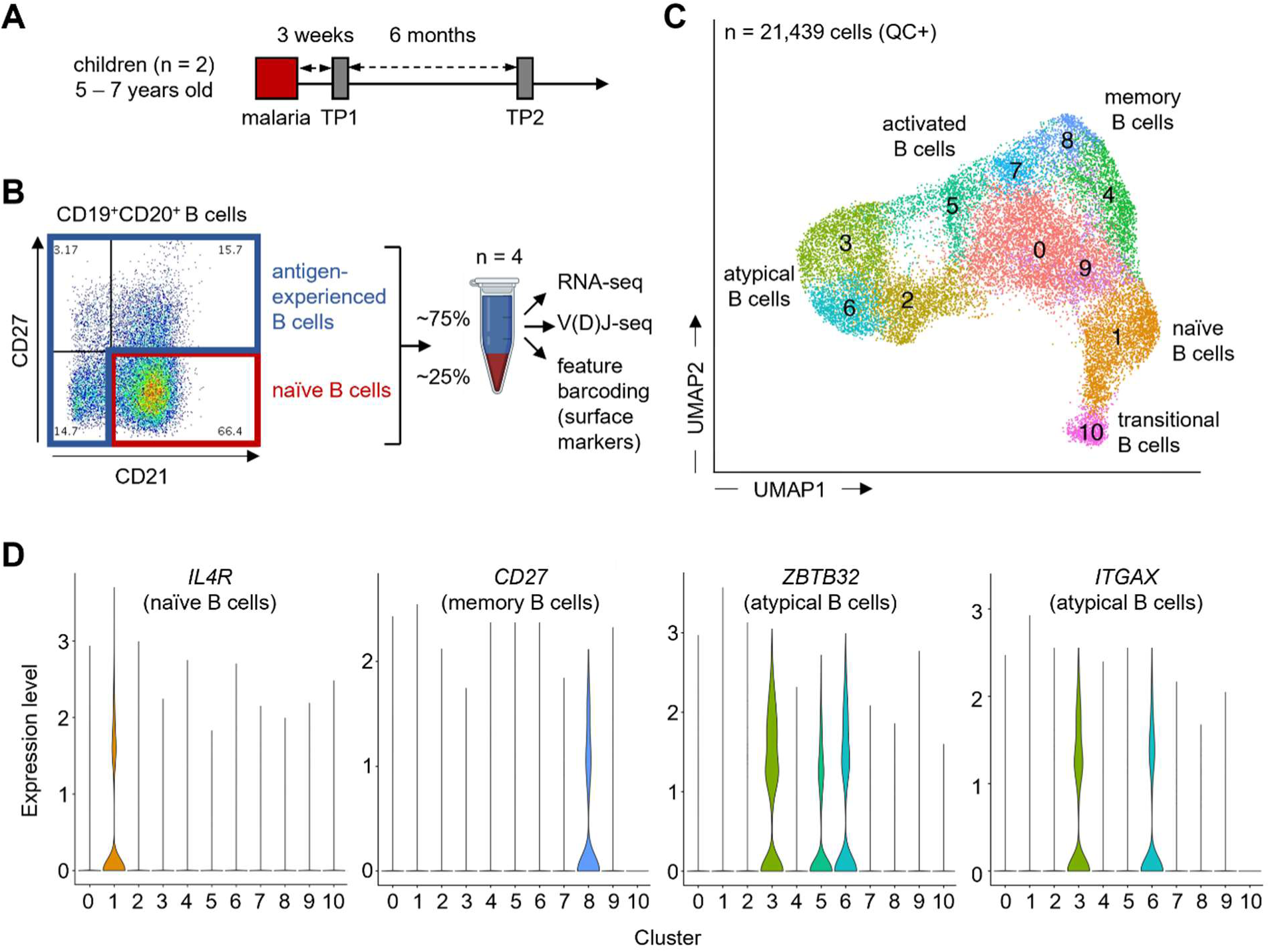
Single-cell sequencing of B cells in malaria-experienced children. **A)** Timeline of sample collection at three weeks post-malaria (TP1) and six months later (TP2) in two children. Children did not experience malaria between the first and second sample collection. **B)** Flow cytometry strategy to enrich for antigen-experienced B cells. A majority of antigen-experienced B cells were mixed with naïve B cells and used for library preparation using the 10x Genomics platform. This yielded three different libraries for each sample to study gene expression, V(D)J sequences, and cell surface marker expression in the same cells. **C)** UMAP of all B cells that passed quality control (QC+), showing 11 transcriptomic B cell clusters. Major subsets identified based on the expression of key genes and previously published gene signatures are indicated. **D)** Expression levels of genes characteristic for naïve B cells (*IL4R*), memory B cells (*CD27*), and atypical B cells (*ZBTB32* and *ITGAX*). TP, time point.

We first isolated naïve and antigen-experienced B cells by fluorescence-activated cell sorting based on the expression of CD21 and CD27 (**Figure 1B**). Naïve B cells typically make up a large fraction of the total B cell pool (43 – 60% in the samples used here), but are not of interest when studying the response of B cells to infection. Therefore, we enriched our samples for antigen-experienced B cells, but added in some naïve B cells to have representation of all B cell populations in the final cell pool. (**Table S2**). Each sample was then subjected to single-cell RNA-sequencing in combination with analysis of select cell surface marker expression and B cell receptor-sequencing on the 10x Genomics platform. Cells that passed quality control (n = 21,439; **Table S3**) were clustered into eleven populations based on gene expression and visualized by Uniform Manifold Approximation and Projection (UMAP) (**Figure 1C**). UMAP visualizes cell clusters in a 2D plot based on similarity in transcriptional profile. Each of the four samples was well represented in all eleven clusters (**Figure S2; Table S4**). Notably, immunoglobulin-related genes were excluded during clustering to prevent antibody isotype or V(D)J-gene segment usage from influencing these results, as was previously proposed by Stewart *et al*. (*20*). Comparing the expression levels of individual key genes (**Figure 1D**), as well as gene signatures of various B cell populations generated by others (**Figure S3**) (*11, 20, 21*), allowed us to identify the location of major B cell populations in the UMAP. This included transitional B cells in cluster 10, naive B cells (expressing *IL4R*) in cluster 1, and memory B cells (expressing *CD27*) in cluster 8. In total, 83% of all cells in our data set were found in clusters with a non-naive transcriptional signature, confirming that our data set was enriched for antigen-experienced cells. Clusters 3, 5, and 6 expressed genes associated with atypical B cells, such as *ZBTB32*, encoding a transcription factor, and *ITGAX,* encoding the surface protein CD11c (**Figure 1D**). Together, these two clusters represented almost 30% of all data (6,088 cells), making this a rich dataset to study heterogeneity among atypical B cells in malaria-experienced individuals.

### Phenotypically defined atypical B cells have multiple distinct transcriptomic profiles

We next sought to determine how well the atypical B cell clusters that we identified based on transcriptomic profiles corresponded to CD21^-^CD27^-^ B cells, which is the phenotypic classification for atypical B cells most commonly used in the malaria field. We therefore used cell surface markers CD21 and CD27 to ‘gate’ cells, similar to how this is commonly done in flow cytometry (**Figure S4**). Nearly all cells in clusters 3 and 6 were CD21^-^CD27^-^, in agreement with their transcriptomic profile of atypical B cells (**Figure 2A, Figure S5**). However, CD21^-^ CD27^-^ B cells also made up about 50% of cluster 5. Cluster 10 was also comprised of approximately 50% CD21^-^CD27^-^ cells, which were transitional B cells that have recently emigrated from the bone marrow and have yet to upregulate CD21 expression (**Figure S3A**). Additionally, cluster 2 largely consisted of CD21^-^CD27^-^ cells that had not undergone class switching (see also below). The CD21^-^CD27^-^ cells in clusters 2 and 10 were therefore excluded from our analysis. In a recent study by Sutton *et al.*, atypical B cells were observed to predominantly express CD11c and CXCR3 (*3*). We therefore also analyzed the distribution of CD11c^+/-^ CXCR3^+/-^ populations among the nine B cell clusters (**Figure S6**). The large majority of cells in clusters 3 and 6 expressed CD11c (75% and 98%, respectively), while expression of CXCR3 was observed in only ∼45% of all cells in these clusters (**Figure 2B, Figure S5**). In contrast, 79% of cells in cluster 5 were CXCR3^+^, most of which also expressed CD11c. These percentages were similar when limiting the analysis to CD21^-^CD27^-^ B cells (**Figure S7**). These data suggest that CXCR3 marks a subset of atypical B cells but cannot be used as a marker to identify all atypical B cells. Next, we analyzed the heavy chain sequences from the B cell receptor-sequencing libraries to determine isotype usage among the different clusters. This analysis showed that, as expected, the transitional cells in cluster 10 and naïve B cells in cluster 1 were unswitched cells, expressing only *IGHM* and *IGHD*. Atypical B cell clusters 3, 5, and 6 contained larger fractions of class-switched B cells (**Figure 2C, Figure S5**). Cluster 3 in particular showed a distribution of *IGH* transcripts approaching that of cluster 8 memory B cells, with the exception of a lack of *IGHA1* expression, suggesting that cluster 3 atypical B cells have undergone class-switch recombination at similar rates as memory B cells.

**Figure 2:**
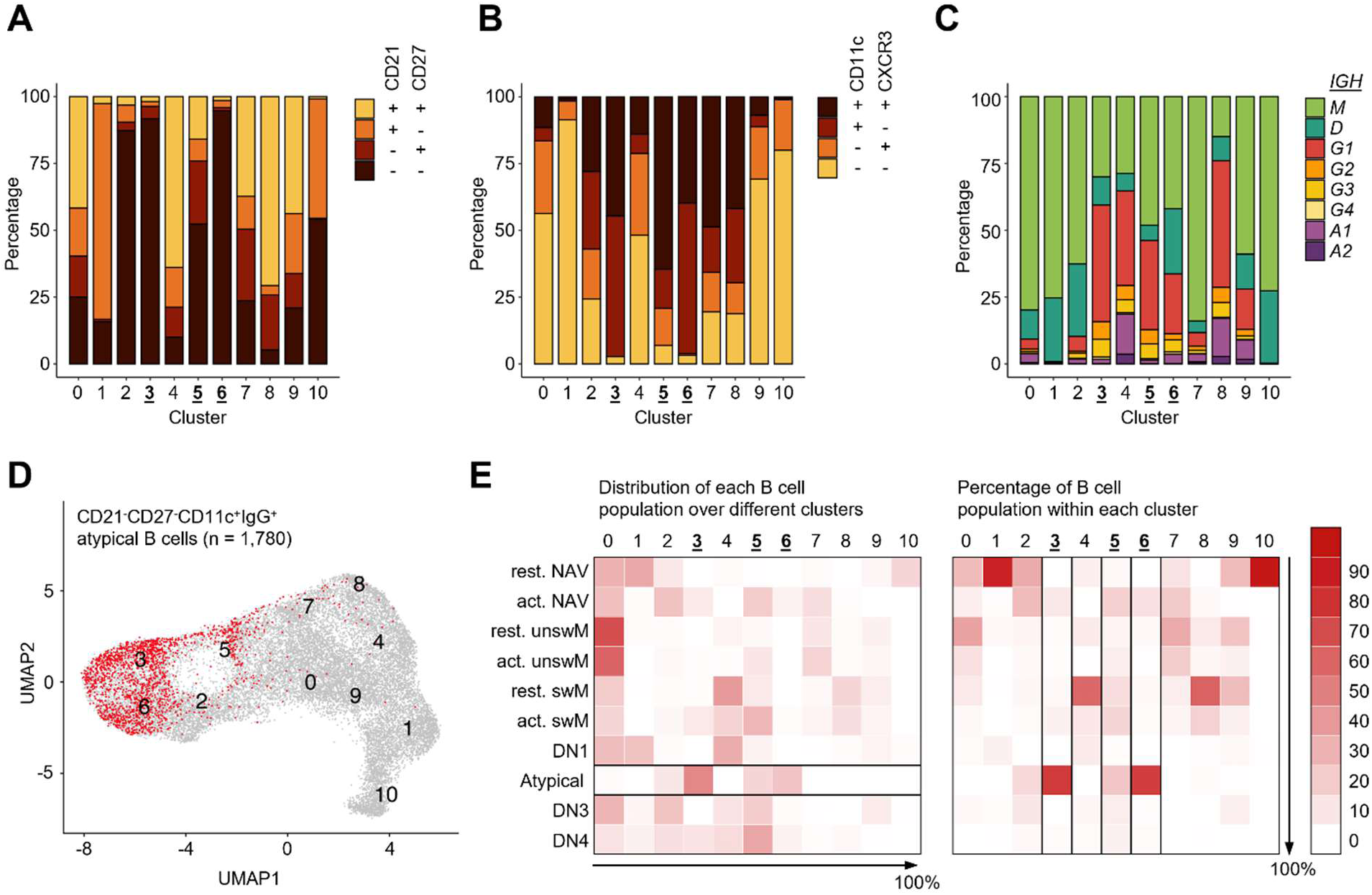
Phenotypically defined atypical B cells are found in three transcriptionally distinct clusters. **A)** Distribution of CD21^-/+^ CD27^-/+^ B cells among all transcriptional clusters. **B)** Distribution of CD11c^-/+^ CXCR3^-/+^ B cells among all transcriptional clusters. **C)** Distribution of immunoglobulin heavy chain transcripts among all transcriptional clusters. **D)** Projection of phenotypically defined atypical B cells (shown in red) onto the transcriptomics-based UMAP. All non-atypical B cells are shown in gray. **E)** The distribution of each B cell population over the different clusters (left) and the percentage of phenotypically defined B cell populations within each cluster (right). The direction in which the values in the rows (left) or columns (right) add up to 100% is indicated with an arrow. The atypical B cell population (left heatmap) and clusters 3, 5, and 6 (right heatmap) mentioned in the text are highlighted with borders. Rest., resting; act., activated; NAV, naïve, unswM, unswitched memory; swM, switched memory; DN, double negative.

To further delineate the distribution of different B cell populations among our transcriptomic clusters, we identified B cells by phenotype based on the surface expression of CD21, CD27, CD11c, IgM, and IgG following recently published guidelines (*22*). Because our cell surface marker analysis did not include IgD, we made two exceptions to these guidelines. First, unswitched B cells were not identified based on the presence of IgD but that of IgM, which is usually co-expressed with IgD. Second, switched B cells were identified based on the presence of IgG instead of the absence of IgD, isotypes that are mutually exclusive. We first used CD27 and IgM or IgG expression to distinguish naive B cells (NAV, CD27^-^IgM^+^), unswitched memory B cells (unswM, CD27^+^IgM^+^), and switched memory B cells (swM, CD27^+^IgG^+^). These populations were further divided into resting (CD21^+^) and activated (CD21^-^). The majority of resting switched memory B cells were located in clusters 0, 4, and 8. We also identified DN B cells (CD27^-^IgG^+^), which were divided into DN1 (CD21^+^CD11c^-^), DN2 (CD21^-^CD11c^+^), DN3 (CD21^-^CD11c^-^), and DN4 (CD21^+^CD11c^+^) (**Figure S8**). IgM^+^ atypical B cells are also known as activated naive B cells, and IgG^+^ atypical B cells are equivalent to DN2 B cells. We recently reported substantial differences in intrinsic properties between IgM^+^ and IgG^+^ atypical B cells that are likely related to their origin and function (*23*). Additionally, CD11c has been identified as the marker that best delineates atypical B cells (*3, 24*). We therefore decided to use the phenotypic definition of DN2 B cells to identify atypical B cells for the remainder of this study: CD21^-^CD27^-^IgG^+^CD11c^+^ (**Figure 2D**). Of all phenotypically defined atypical B cells, 47% were found in cluster 3, 23% in cluster 6, and 17% in cluster 5 (**Figure 2E**). The majority of cells in cluster 3 (77%) and cluster 6 (75%) were atypical B cells (**Figure 2E**). Cluster 5 was more diverse with 23% atypical B cells, 19% activated naïve B cells, 13% resting switched memory B cells, 16% activated switched memory B cells, and 29% other B cell populations. Of note, clusters 3, 5, and 6 were stable at a clustering resolution ≥ 0.6. At a clustering resolution below 0.6, cluster 6 was merged with cluster 2 that predominantly contained activated naïve B cells (**Figure S9**). For this reason, a resolution of 0.6 was chosen for the analysis presented here. Collectively, these results show that phenotypically defined class-switched atypical B cells in malaria-experienced children consist of multiple distinct subsets with unique transcriptional profiles.

### Differences in gene expression programs of atypical B cells are associated with different roles in the immune response

To better understand the identity of the three subsets of atypical B cells identified in this study found in clusters 3, 5, and 6, we compared the expression of genes that have previously been described as part of the unique transcriptional profile of atypical B cells (*10–12*). The up or down regulation of various genes in these clusters as compared to resting switched memory B cells in cluster 8 was consistent with previous reports (**Figure 3A**). Two manuscripts recently posted on bioRxiv showed that the transcription factor *ZEB2* acts as an important regulator for the development of atypical B cells (*25, 26*). ZEB2 was also expressed significantly higher in clusters 3, 5, and 6 as compared to cluster 8 (**Figure 3A**). In contrast, several atypical B cell-associated genes were uniquely expressed in only one of the three clusters (**Figure 3B**). For example, the integrin-encoding genes *ITGB2* and *ITGB7* were predominantly expressed in cluster 3. In contrast, *CD72*, encoding a surface marker that inhibits B cell responses and prevents antibody production (*27*), was more abundantly expressed in cluster 6. *CXCR3* expression was highest in cluster 5, in line with the results of the cell surface marker analysis (**Figure 2B**). These differences in gene expression highlight that bulk atypical B cells are a mixture of various subsets and point to functional differences in activation status and the potential for cell-cell interactions between these subsets.

**Figure 3:**
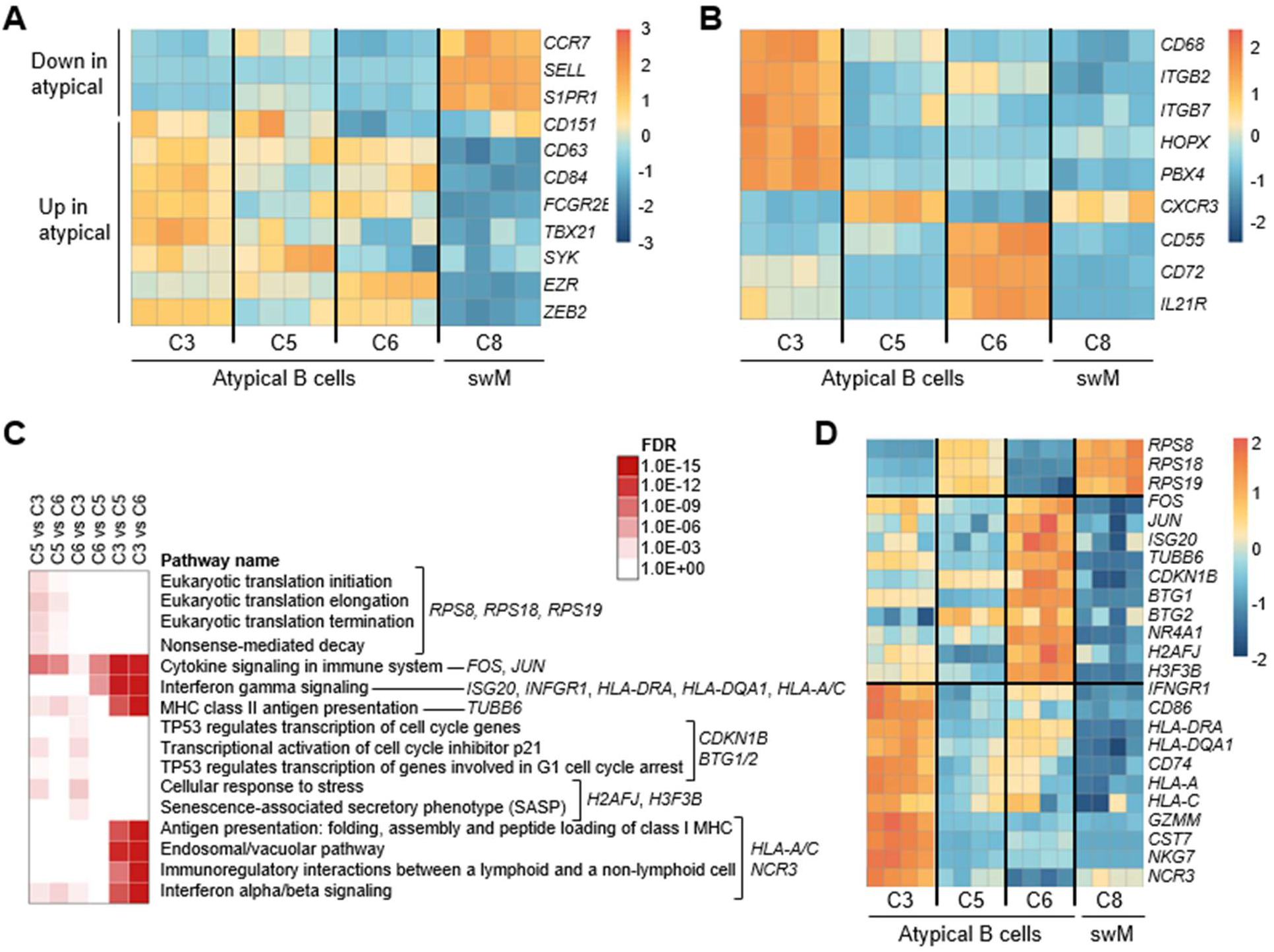
Phenotypic and functional differences between subsets of atypical B cells. **A)** Expression of genes previously identified as part of the unique transcriptional signature of atypical B cells that are consistently expressed higher or lower in all three subsets of atypical B cells as compared to class-switched resting memory B cells (swM; cluster 3). **B)** Differential expression of atypical B cell-associated genes between the three subsets of atypical B cells. **C)** Selected reactome pathways that were significantly enriched (FDR-corrected P value < 10^-4^) in pair-wise comparisons between the three clusters that contain atypical B cells. Key differentially expressed genes in these pathways that are shown in panel D are indicated. **D)** Select differentially expressed genes in atypical B cell clusters that were present in enriched Reactome pathways and are discussed in the text. The four columns represent the four samples subjected to scRNA-seq analysis. C8, cluster 8; C3, cluster 3; C5, cluster 5; C6, cluster 6

To gain insight into potential differences in function between cells in clusters 3, 5, and 6, we performed Reactome pathway analysis using genes that were upregulated or downregulated in pair-wise comparisons of the three clusters (**Figure 3C-D, Tables S5 – S6**). Atypical B cells in cluster 6 showed higher expression of 8 out of 99 genes encoding ribosomal proteins (for example, *RPS8*, *RPS18*, and *RPS19*) and were most strongly enriched for pathways related to translation. Ribosomal genes were also highly expressed in cluster 8 switched memory B cells, suggesting that this may be a feature of cells poised to undergo cell differentiation. Atypical B cells in both clusters 3 and 6 upregulated genes in response to stimulation by interferon gamma and cytokines, but the outcome of this response differed between the two clusters. Atypical B cells in cluster 6 upregulated genes encoding components of the Activator Protein 1 (AP1) complex, including *FOS* and *JUN*, as well as interferon-stimulated genes (for example, *ISG20*), and tubulin genes (for example, *TUBB6*). Additionally, cluster 6 atypical B cells were enriched for pathways that signified negative regulation of cell cycle progression. Upregulated genes included the cyclin-dependent kinase inhibitor *CDKN1B*, the anti-proliferative genes *BTG1/2*, and the transcriptional repressor *NR4A1*. Nuclear receptor 4A1 is part of a family of transcription factors that was recently implicated in a negative feedback loop that rendered B cells more dependent on T cell help and promoted peripheral B cell tolerance (*28, 29*). Finally, cluster 6 atypical B cells showed increased expression of variant histones, associated with a pro-inflammatory stress response resulting in cellular senescence (*30*)(*31, 32*). The transcriptional profile of atypical B cells in cluster 6 thus pointed to an interferon-driven response that resulted in senescence or anergy. On the other hand, atypical B cells in cluster 3 were enriched for pathways related to antigen processing and presentation, and immunoregulation. Among the genes upregulated in cluster 3 were genes encoding proteins that promote interactions between B cells and T cells, such as HLA-DR/DQ, CD86, which is downregulated by NR4A1, and CD74, which regulates antigen presentation by class II MHC (*33–35*). In addition, we observed upregulation of genes encoding class I MHC (*HLA-A* and *HLA-C*) and several genes that are associated with cytotoxicity, including granzyme M (*GZMM*), cystatin 7 (*CST7*), natural killer cell granule protein 7 (*NKG7*, a target gene of T-bet), and natural cytotoxicity triggering receptor 3 (*NCR3*). This gene signature suggests that these cells have increased exocytosis activity, which could affect cell-cell interactions and have an immunomodulatory function. Collectively, our results indicate that phenotypically defined IgG^+^ atypical B cells can have one of three different transcriptional profiles that are associated with different functions within the immune response.

### Atypical B cells form two separate compartments

To study the development of the different subsets of atypical B cells, we turned to an analysis of their B cell receptor sequences. We first assessed the amount of affinity maturation each B cell population has undergone by determining the level of somatic hypermutation in each cluster. This revealed that clusters 3 and 5 on average accumulated similar levels of mutations as class-switched memory B cells in cluster 8, while the average mutation frequency in cluster 6 was more than 50% lower (**Figure 4A, Figure S10A**). However, this may in part be caused by the larger fractions of unswitched B cells in cluster 6 (**Figure 2C**). When including only class-switched atypical B cells, we still observed a higher average level of somatic hypermutation in clusters 3 and 5 than in cluster 6, although the difference was slightly smaller (**Figure 4B, Figure S10B**). This difference suggests that cluster 6 atypical B cells have not undergone extensive affinity maturation and seem to be more closely related to naïve B cells. Atypical B cells in clusters 3 and 5 have undergone similar levels of affinity maturation as memory B cells, which could indicate that they are derived from memory B cells or have a similar developmental trajectory. However, while somatic hypermutation and affinity maturation typically occur inside germinal centers, these processes can also occur in extrafollicular sites (*36*) and our results are therefore not direct evidence for the developmental pathway of atypical B cells.

**Figure 4:**
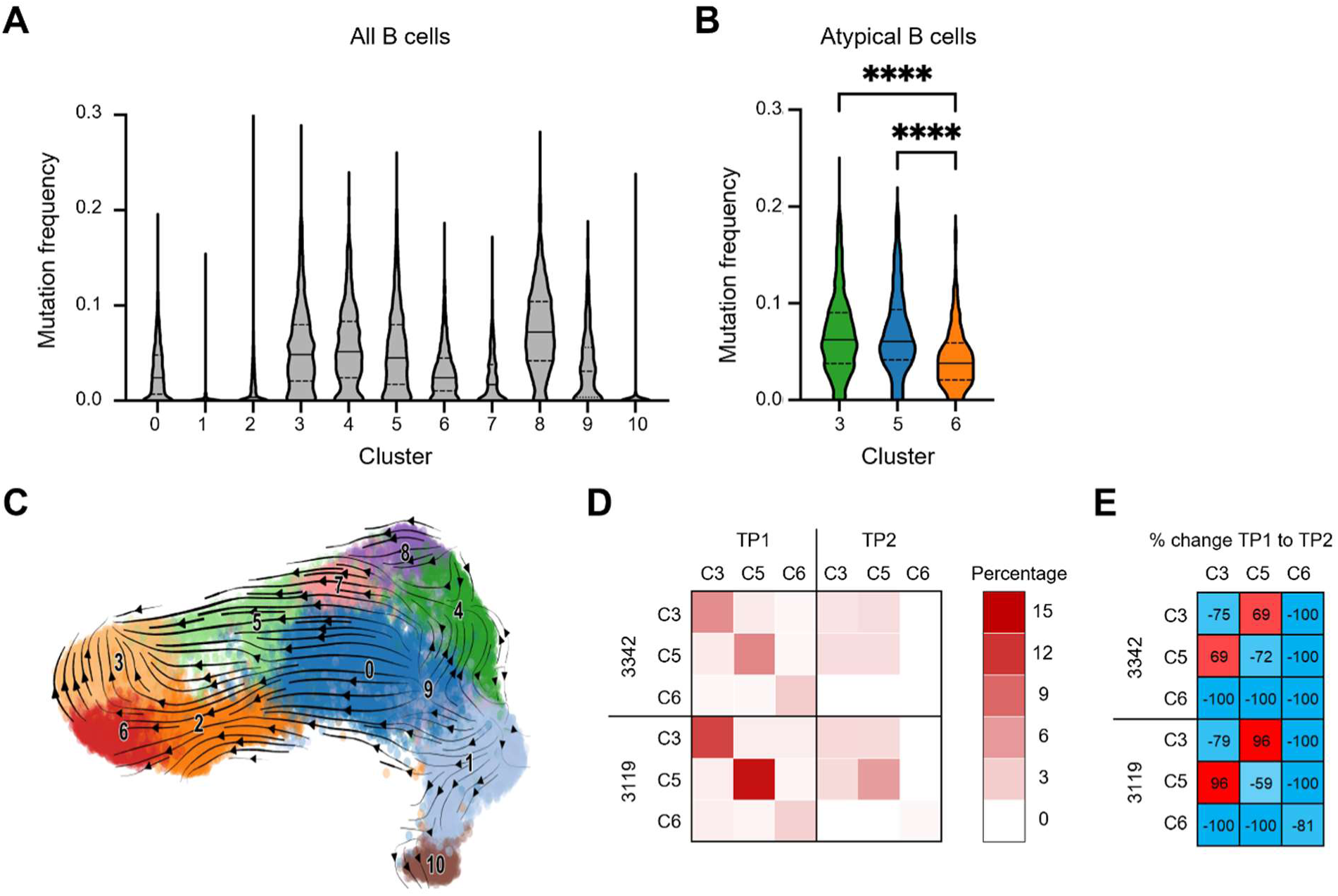
Developmental relationships between atypical B cell subsets. **A)** Levels of somatic hypermutation among the transcriptomics-based B cell clusters. **B)** Levels of somatic hypermutation among phenotypically defined atypical B cells in clusters 3, 5, and 6. In panels A and B, the median is shown as a solid horizontal lines, while the quartiles are represented by dashed horizontal lines. In panel B, a one-way ANOVA was used to test for statistically significant differences between the groups. P values shown are from Tukey’s post hoc test. **** P < 0.0001. **C)** Analysis of the developmental trajectory of B cells using scVelo. **D)** Clonal expansion within clusters (along the diagonal in each quadrant of the heatmap) and clonal connections between clusters (off-diagonal cells) for the three clusters that contain atypical B cells at 3 weeks post-malaria (TP1) and six months later (TP2). **E)** The percentage change in clonal expansion/connections between the first and the second time point for the three clusters that contain atypical B cells. TP, time point

Next, we wished to better understand the developmental relationship between the three subsets. To this end, we first employed RNA velocity analysis to infer the developmental trajectories of the three atypical B cell subsets. This analysis uses the ratio of unspliced and spliced transcripts to identify genes that undergo upregulation (more unspliced than spliced transcripts) or downregulation (more spliced than unspliced transcripts). These transcript dynamics are then used to model the direction and speed of movement of individual cells (*37*). This analysis revealed two dominant developmental trajectories: (1) cluster 6 atypical B cells were mainly derived from activated naïve B cells in cluster 2 and did not develop further, while (2) cluster 3 atypical B cells seemed to be predominantly derived from memory B cells, with cluster 5 atypical B cells as an intermediate step in this trajectory (**Figure 4C**). To confirm these developmental connections between the atypical B cell subsets, we calculated the level of expansion of clonal B cell lineages within each cluster at the two time points, as well as the percentage of clonal connections between clusters, as described in more detail previously (*38*). B cells were considered part of the same lineage when their heavy chain variable region contained the same *IGHV* and *IGHJ* genes, and shared at least 85% amino acid sequence identity in the heavy chain (H) CDR3 (*39, 40*). In all three atypical B cell clusters, there was strong expansion of B cells at 3 weeks post-malaria, with 3 – 14% of B cells within each cluster belonging to the same lineage (**Figure 4D, Figure S11A**). This was followed by contraction six months later, resulting in a loss of more than 50% of clonal lineages (**Figure 4D, E**). This contraction points to a return to a resting state and suggests that minimal immune activation occurred as a result of the asymptomatic infection at 6-months post-malaria, as compared to the strong immune response elicited by malaria. At 3 weeks post-malaria, 0.5 – 1.0% of clonal sharing occurred between cluster 6 and clusters 3/5, but these connections were completely lost six months later. In contrast, the connectivity between cluster 3 and cluster 5 increased at the second time point, as compared to the time point shortly after malaria, and this was the only pair of clusters for which this happened consistently (**Figure 4E, Figure S11B**). These observations suggest that cluster 6 atypical B cells form a separate compartment from the other atypical B cell subsets, while cluster 3 and cluster 5 atypical B cells are more strongly connected, in line with the results of the trajectory analysis. Collectively, these results point to different developmental pathways for atypical B cells that are connected to different functional roles in the immune response.

### All three atypical B cell subsets are found in malaria-experienced children and adults irrespective of parasite exposure levels

The identification of atypical B cell clusters by transcriptomics analysis was based on four samples from two children. To validate our findings in a larger number of malaria-susceptible children and extend our analysis to malaria-protected adults, we turned to an analysis of these cells by flow cytometry. In the single-cell transcriptomics analysis, we observed that the three clusters with atypical B cells could be distinguished by differences in expression level of *ITGAX* (encoding CD11c) and *CD86* (**Figure 5A**). We therefore sorted three subsets of atypical B cells based on expression of CD11c and CD86 (**Figure 5B, Figure S12**) and used bulk RNA-seq analysis to confirm that these subsets have similar gene expression profiles as the three transcriptomics clusters identified by scRNA-seq (**Figure 5C**).

**Figure 5:**
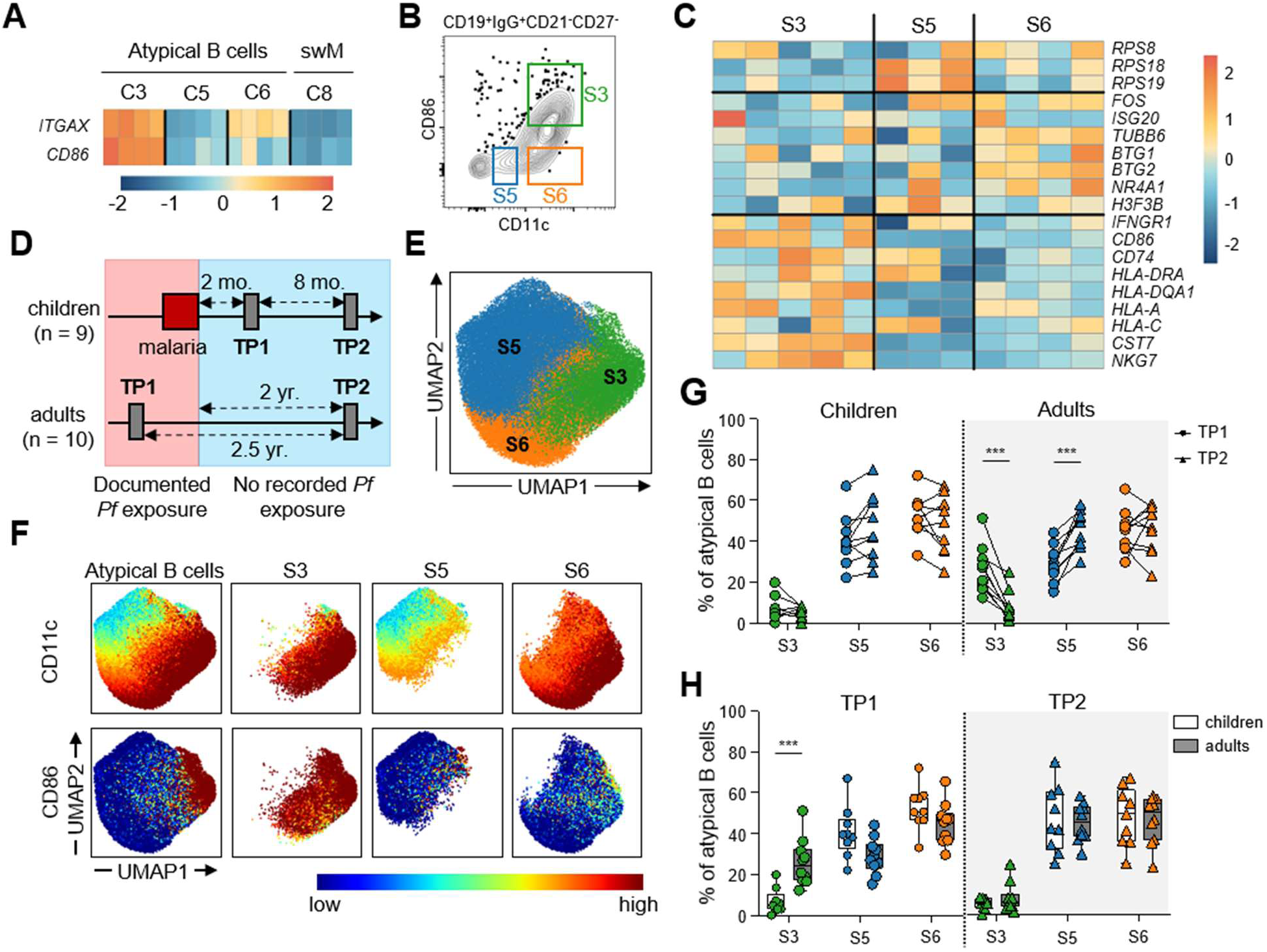
Dynamics of atypical B cell subsets in children and adults. **A)** Gene expression profiles of differentially expressed genes encoding cell surface proteins CD11c and CD86 in atypical B cell clusters 3, 5, and 6, as well as resting switched memory B cells (swM, cluster 8), used to distinguish atypical B cell subsets by flow cytometry. **B)** Gating strategy to sort atypical B cell subsets based on CD11c and CD86 expression. **C)** Expression of genes that underlie the different functional profiles of the three atypical B cell clusters (compare with Figure 3D) in flow-sorted atypical B cells subsets. **D)** Schematic overview of sample collection for children and adults during known *P. falciparum* (*Pf*) exposure and in the absence of exposure to *P. falciparum*. **E)** Composite UMAP projection of atypical B cells from all 38 samples. Subsets were defined based on expression of CD11c and CD86 and were named according to the corresponding clusters defined by single-cell RNA-sequencing analysis. **F)** CD11c and CD86 expression projected onto the UMAP of all atypical B cells as well as each of the three subsets individually. **G)** Relative size of each atypical B cell subset in children (n = 9) and adults (n = 10) at the first time point (TP1; circles) and second time point (TP2; triangles). **H)** Comparison of relative subset sizes between children and adults at each time point. Statistical analysis was performed using a Wilcoxon matched-pairs signed rank test for paired data (E) or a Mann-Whitney test for unpaired data (F) with a 10% false discovery rate using the two-stage step-up method of Benjamini, Krieger, and Yekutieli. * P < 0.05; ** P < 0.01; *** P < 0.001.

Using spectral flow cytometry, we next aimed to study the abundance of each atypical B cell subset and their longevity in the absence of antigen stimulation and infection-driven inflammation. To this end, we included samples from nine children and ten adults collected following known *Plasmodium* exposure and months to years later after a period with low *Plasmodium* exposure due to highly effective insecticide spaying (*41, 42*) (**Figure 5D, Table S1**). B cells were phenotyped using a panel of 18 surface and intracellular markers that would allow us to identify the three atypical B cell subsets and analyze these for the expression of markers previously shown to be positively (CD19, CD20, CD11c, FcRL5, CD83, CD86, CD95, CXCR3, T-bet) or negatively (CD24, CD38, CXCR5) associated with activated and atypical B cells (*22*). To find the atypical B cell subsets identified in our transcriptomic analysis, we first gated manually on CD19^+^CD20^+^CD21^-^CD27^-^IgG^+^CD11c^+^ atypical B cells (**Figure S13**). The atypical B cells from all samples were then plotted in a composite UMAP based on the expression of the 12 cell surface and intracellular markers previously shown to be associated with activated and atypical B cells (**Figure S14**).

Next, we used the unsupervised clustering tool FlowSOM to identify three distinct atypical B cell subsets based on the expression of CD11c and CD86 (**Figure 5E**). Projection of these subsets onto the UMAP showed that the expression of *ITGAX* and *CD86* matched the cell surface expression signatures of subset 3 (CD11c^hi^CD86^+^), subset 5 (CD11c^int^CD86^-^), and subset 6 (CD11c^hi^CD86^-^), (**Figure 5F**). We will refer to these subsets with the number corresponding to the matching transcriptomics cluster, i.e., flow cytometry subset 3 for cells matching transcriptomics cluster 3. We then determined the relative abundance of these three subsets among all atypical B cells. In children, most atypical B cells belonged to subset 6 (∼50%) and subset 5 (∼40%). The relative abundance of the subsets did not change between time points (**Figure 5G**, **Figure S15**). This could be the result of the timing of sample collection. At two months post-malaria in a low transmission setting, transient changes to the atypical B cell population induced by malaria may have already reverted to baseline. In adults, the first sample was collected during a period of high *Plasmodium* exposure. Interestingly, the proportion of subset 3 atypical B cells was larger in adults than in children at the first time point but decreased to the level seen in children after a period of minimal parasite exposure (**Figure 5H**). This was also true when expressing the abundance of subset 3 atypical B cells as a percentage of total B cells (**Figure S15**). Simultaneously, we observed an increase in the relative proportion of subset 5 among all atypical B cells between the two time points in the adults, although the percentage of subset 5 atypical B cells among total B cells did not change (**Figure 5G**, **Figure S15**). Collectively, these results demonstrate that we were able to distinguish three subsets of atypical B cells by flow cytometry and that the composition of the atypical B cell population changes in relation to parasite exposure.

### Atypical B cells express higher levels of CXCR3 and CD95 in adults as compared to children

To map additional differences in the phenotype between the three subsets of atypical B cells, we plotted the mean fluorescence intensity of CD21, CD27, and the 12 markers that were previously observed to be positively or negatively associated with activated or atypical B cells for all three subsets as well as IgG^+^ resting memory B cells (CD21^-^CD27^-^). We observed that subset 3 had the most ‘atypical’ surface marker expression pattern, with the highest expression of CD19, CD20, CD11c, FcRL5, CD95, and T-bet (**Figure 6A**, **Figure S16**).

**Figure 6:**
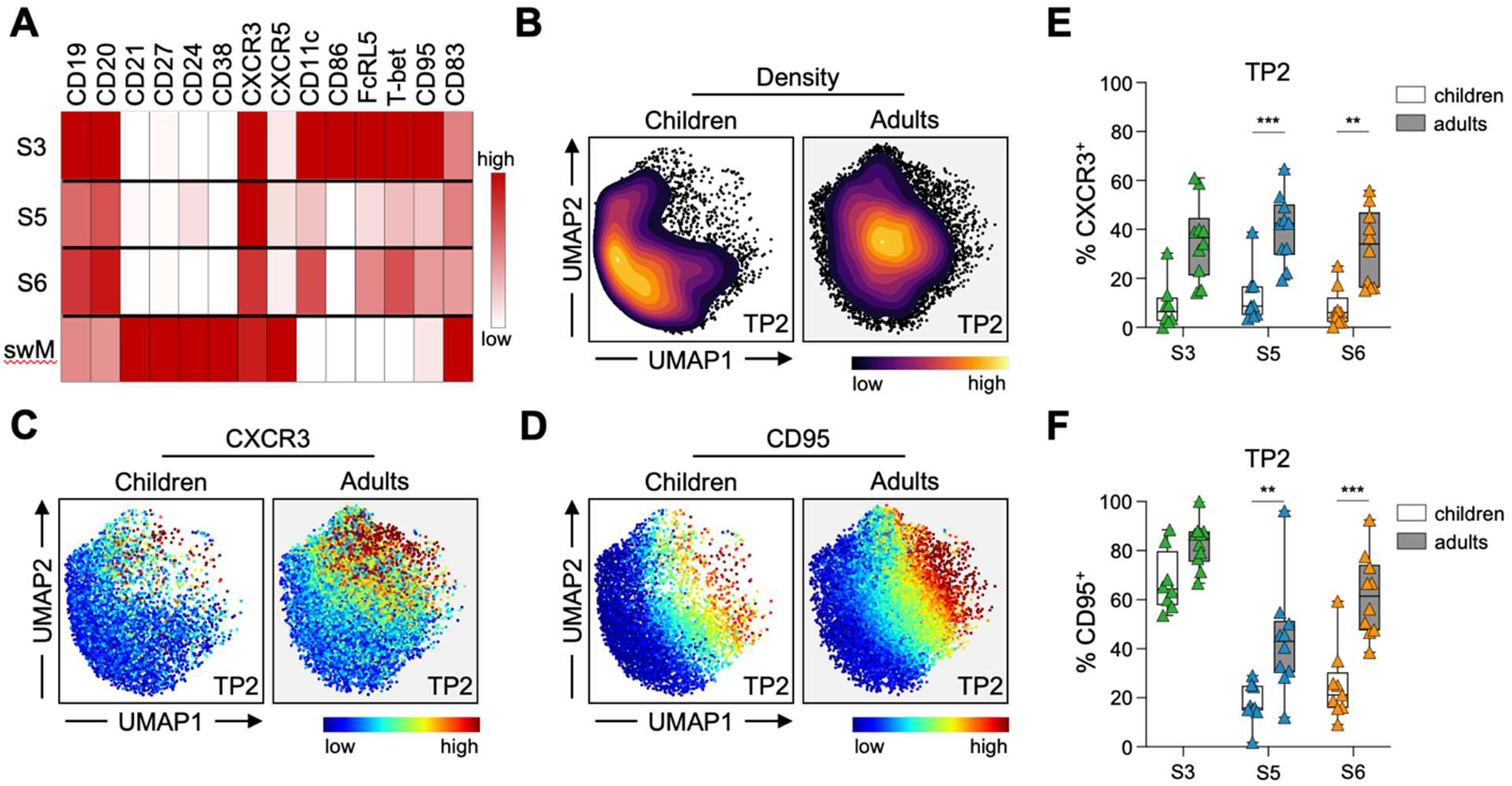
Expression of T-bet and CXCR3 in atypical B cell subsets. **A)** Relative median fluorescence intesity of 14 surface and intracellular markers in subsets (S) 3, 5, and 6 and in resting switched memory B cells (swM). **B)** Density UMAP plots of atypical B cells detected at the second time point in children and adults. **C-D)** Projection of CXCR3 (C) and CD95 (D) expression in atypical B cells at the second time point projected onto the UMAPs for children and adults. **E-F)** Percentage of CXCR3^+^ (E) and CD95^+^ (F) cells within atypical B cell subsets. For each subset, only individuals with >50 cells at both time points were included in the analysis. Statistical analysis was performed using a Mann-Whitney test with a 10% false discovery rate using the two-stage step-up method of Benjamini, Krieger, and Yekutieli. * P < 0.05; ** P < 0.01; *** P < 0.001.

We were also interested in potential differences in phenotype between atypical B cells from malaria-experienced children and adults. At the second time point, after months to years of no chronic immune activation, the distribution of atypical B cell subsets was similar between children and adults (**Figure 5H**). However, we noticed that their atypical B cells had a distinct localization when projected onto the UMAP (**Figure 6B**). This observation suggests that atypical B cells from children and adults differ in the expression of one or more other markers. Overlaying the expression of other markers in our flow cytometry panel onto the UMAP revealed that atypical B cells in adults extended into the UMAP area where cells expressed high levels of CXCR3 and CD95 (**Figure 6C, D**). To quantify this, we analyzed the percentage of CXCR3^+^ and CD95^+^ cells in each subset. Adults showed more CXCR3^+^ cells in all atypical B cell subsets as compared to children (**Figure 6E**), and more CD95^+^ cells in subsets 5 and 6 (**Figure 6F**). Collectively, this difference in CXCR3 and CD95 expression between children and adults points towards additional heterogeneity among atypical B cells that may be connected to differences in immune status or parasite exposure levels.

### All subsets of atypical B cells harbor *P. falciparum*-specific cells

To determine whether the different atypical B cell subsets develop in response to antigen during infection or through bystander activation, we used antigen tetramers to detect B cells with specificity to two *P. falciparum* proteins: merozoite surface protein 1 (MSP1) and apical membrane antigen 1 (AMA1). Both antigens are known to elicit strong IgG responses in children and adults (*43*). Since both MSP1 and AMA1 tetramers were labeled with the same fluorochromes, MSP1-specific and AMA1-specific B cells are together referred to as MSP1/AMA1-specific (**Figure S13**). First, we overlayed the MSP1/AMA1-specific B cells onto the previously generated UMAPs of subset 3, 5, and 6 atypical B cells. Antigen-specific cells were identified in all subsets (**Figure 7A**). We then quantified the number of antigen-specific cells in each of the three atypical B cell subsets and observed that the distribution of MSP1/AMA1 B cells over the three subsets mirrored their total size (**Figure 7B**). Finally, we analyzed CXCR3 and CD95 expression among MSP1/AMA1-specific atypical B cells in adults. Approximately 25 - 40% of all MSP1/AMA1-specific atypical B cells were CXCR3^+^, with no clear pattern related to subset identity (**Figure 7C**). In contrast, nearly all MPS1/AMA1-specific atypical B cells in all three subsets expressed CD95, which was an increase as compared to all atypical B cells in subsets 5 and 6 (**Figure 7D**). Increased CD95 expression was also observed in Spike-specific B cells in recovered COVID-19 patients (*44*) and suggests that these cells recently underwent antigen-driven activation. Collectively, these results suggest that all three subsets of atypical B cells develop in response to antigen stimulation.

**Figure 7:**
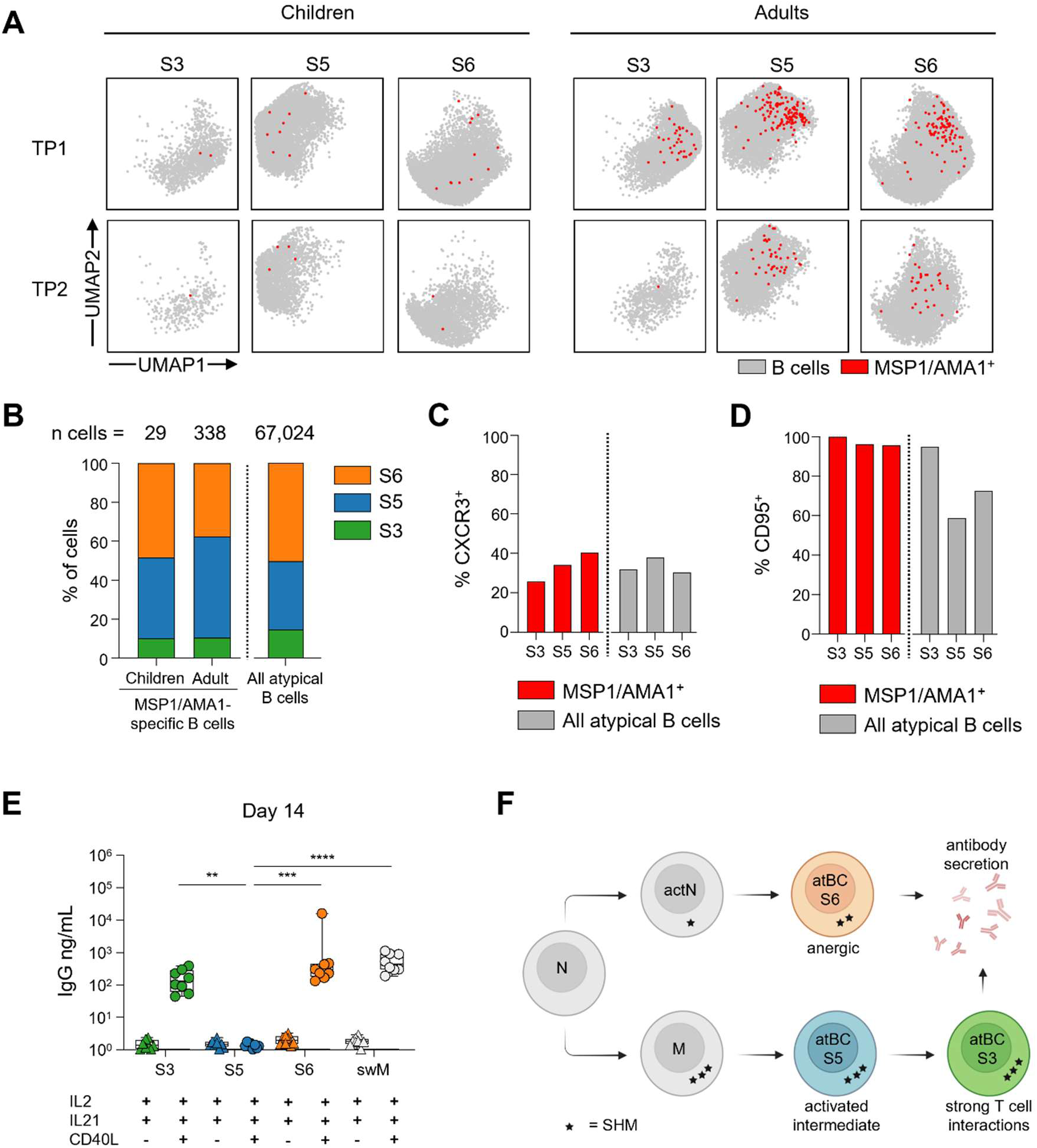
Antigen-specificity and antibody secretion among the three atypical B cell subsets. **A)** Projection of MSP1/AMA1-specific B cells (red) onto the UMAPs of atypical B cell subsets, shown separately for time point 1 (TP1) and time point 2 (TP2) in 9 children and 10 adults. **B)** Distribution of all MSP1/AMA1-specific B cells over the three atypical B cell subsets in children and adults. Data from both time points were combined. **C)** Percentage of all CXCR3^+^ antigen-specific cells in each atypical B cell subset for adult samples. **D)** Percentage of all CD95^+^ antigen-specific cells in each atypical B cell subset for adult samples. **E)** IgG concentration in culture supernatant after 14 days of in vitro culture. A one-way ANOVA was used to test for statistically significant differences between the groups. P values shown are from Kruskal-Wallis post hoc test. ** P < 0.01; *** P < 0.001; **** P < 0.0001. **F)** Proposed model of the effector functions and developmental pathways of atypical B cell subsets.

### Differences in the level of antibody secretion in atypical B cell subsets receiving T cell help

Recently, Hopp et al. observed that bulk atypical B cells express higher levels of markers associated with T cell interactions than memory B cells (*17*). With the help of activated T follicular helper cells, atypical B cells could be stimulated to secrete antibodies, although antibody concentrations were much lower than for memory B cells (*17*). Based on our observations that only two out of the three atypical B cell subsets expressed markers involved in T cell interactions (S3 and S6), we sought to determine which atypical B cell subsets can be induced to secrete antibody under T-cell dependent activation. To this end, we FACS-sorted the three atypical B cell subset based on CD11c and CD86 expression, as well as switched memory B cells from eight malaria-experienced adults (**Figure S12**). We then cultured each B cell population with CD40L-expressing 3T3 cells supplemented with IL-2 and IL-21 to mimic T-cell dependent stimulation, and measured IgG production after 7 and 14 days. The same culture conditions without CD40L-expressing 3T3 cells were included as a negative control. The distribution of atypical B cells over the three subsets was reflective of high *P. falciparum* exposure, allowing us to interrogate their function in terms of recent antigen exposure (**Figure S17A**). We found that both subset 3 and subset 6 secreted IgG in the presence of T-cell help. On day 7, the IgG concentration was lower than that produced by switched memory B cells (**Figure S17B**), but this difference disappeared by day 14 (**Figure 7E**). Subset 5 atypical B cells secreted only minimal amounts of IgG under these conditions. These results suggest that most but not all atypical B cells can differentiate into antibody-secreting cells upon T-cell dependent activation.

## DISCUSSION

Atypical B cells accumulate in a range of conditions that share chronic immune activation as the common denominator, including chronic or repetitive infection and autoimmunity. Attempts to understand the function of these cells have resulted in conflicting observations. We hypothesized that these different observations are caused by heterogeneity among atypical B cells and that the underlying subsets are indistinguishable with the markers conventionally used to define atypical B cells. Here, we used multimodal single-cell measurements to better define the heterogeneity among atypical B cells using samples from individuals with chronic *Plasmodium* exposure. We identified novel subsets of atypical B cells, each with a unique transcriptional program and cell surface marker signature, that are associated with different effector functions in the host immune response. Using spectral flow cytometry, we confirmed the presence of these atypical B cell subsets in a cohort of malaria-experienced children and adults, determined that they can develop in response to antigen stimulation, and assessed their ability to differentiate into antibody-secreting cells.

We generated single-cell sequencing data for almost 21,500 B cells, of which we were able to assign slightly more than 14,000 cells to one of 10 phenotypically defined B cell populations. Among these were just over 4,000 cells that belonged to three clusters containing phenotypically defined atypical B cells. Two other recent studies have attempted to assess potential heterogeneity among atypical B cells using single-cell transcriptomics approaches. Holla *et al*. subclustered 199 B cells from malaria-experienced individuals with an atypical transcriptomics profile and observed three subsets that were mainly discriminated by isotype usage (i.e., IgD^+^IgM^+^, IgD^+^IgM^lo^, and IgD^-^IgG^+^ atypical B cells) (*11*). Too few cells were present in that data set to observe subclusters within the IgD^-^IgG^+^ atypical B cell group, which was the focus of our study. Sutton *et al.* analyzed a larger number of IgD^-^ B cells (7,167 cells from malaria-experienced individuals and 5,813 cells from individuals not exposed to *Plasmodium* infection) that were divided into three atypical B cell clusters and several other B cell clusters (*3*). However, reanalysis of this data set did not reveal a clear separation into the same subsets of atypical B cells that we observed in our data set. The main reason for this observation could be that Sutton *et al.* used samples from immune adults who had not recently experienced a malaria episode. Likewise, the gene expression profiles of atypical B cell subsets 3 and 6 from malaria-experienced adults that were generated in this study were very similar (**Figure S18**), although differences in expression of genes that underlie the different function profiles of the three atypical B cells subsets were still apparent (**Figure S5C**). A malaria episode may drive these functionally distinct gene expression programs, followed by their convergence in the absence of such a strong stimulus.

Using our single-cell transcriptomics data set of atypical B cells, we confirmed similarities in gene expression between our three atypical B cell clusters and bulk RNA sequencing transcriptional programs that have previously been reported for atypical B cells (*10, 12*). However, we also discovered that some genes previously linked to atypical B cells were expressed in only one of these three clusters and may be connected to the different functional profiles of these cells. For example, CD72 negatively regulates B cell activation (*27*). *CD72* was uniquely expressed in cluster 6 that has a profile of senescence or anergy. Cluster 3, on the other hand, expressed higher levels of the gene encoding CD68, a transmembrane glycoprotein that is mainly associated with the endosomal/lysosomal compartment in monocytes and macrophages, but is also expressed by lymphocytes (*45, 46*). This observation may be related to the pathways of antigen processing and exocytosis that were upregulated in this cluster.

Using differentially expressed surface markers, we were able to identify three corresponding subsets of atypical B cells by flow cytometry. All three subsets harbored *P. falciparum*-specific B cells, suggesting that each of these subsets can develop as part of a specific response to pathogen-derived antigens and not merely through bystander activation. The lower levels of somatic hypermutation set cluster 6 apart from clusters 3 and 5. Indeed, RNA velocity and B cell receptor repertoire analyses suggested that cluster 6 formed a compartment that was mostly separated from clusters 3 and 5, and followed a different developmental trajectory. The RNA velocity analysis, the increase in percentage of B cell clones that were shared between clusters 3 and 5 over time as the B cell response contracted to baseline levels, and the observation that the decrease in subset 3 atypical B cells over time was paralleled by a relative increase in the size of subset 5 together suggest that atypical B cells in subsets 3 and 5 are developmentally related.

Based on these data, we propose the following working model (**Figure 7F**). Chronic immune activation results in the development of three subsets of atypical B cells. Subset 6 is derived from activated naïve B cells and has relatively low levels of somatic hypermutation. These characteristics are similar to atypical B cells in SLE patients that arise through extrafollicular activation. Under conditions that mimic the interaction with CD4^+^ T cells, subset 6 atypical B cells can further differentiate into short-lived antibody-secreting cells, although these cells are likely to have a high activation threshold. The main developmental trajectory of subset 3 atypical B cells is from memory B cells via an activated subset 5 state. These cells have undergone high levels of somatic hypermutation, consistent with the normal development of memory B cells. Subset 3 atypical B cells are primed for interactions with T cells and will differentiate into antibody-secreting cells upon stimulation by T cells (*47*). Additionally, these cells express proteins that may modulate the function of T cells in return. Subset 5 atypical B cells express lower levels of markers typically associated with atypical B cells, such as FcRL5, T-bet and CD11c, and do not differentiate into by interaction with T cells alone. Other stimuli, such as BCR engagement, additional cytokines, or TLR engagement may drive the differentiation of this subset into antibody-secreting cells. Indeed, Sullivan *et al*. reported that a subset of atypical B cells that expressed lower levels of FcRL5 showed more antibody secretion upon BCR stimulation than FcRL5^+^ atypical B cells (*10*). Collectively, the functional profiles of the three clusters of atypical B cells recapitulate many of the observations that have been made about the atypical B cell population as a whole. Our results indicate that specific subsets of atypical B cells are likely responsible for conflicting observations related to anergy/tolerance, immune-modulation, and the ability to differentiate into antibody-secreting cells.

Comparing atypical B cells between malaria-experienced children and adults revealed additional heterogeneity. Adults showed a higher percentage of CXCR3^+^ cells across all three subsets of atypical B cells, which did not decrease after a two-year period of minimal *P. falciparum* exposure. Hopp *et al.* reported that expression of *CXCR3* is upregulated in atypical B cells in response to malaria in children (*17*). The stable expression of CXCR3 in adults suggests that this marker is acquired as a result of chronic antigen exposure and may play an important role in the protective immune response against malaria. Sutton *et al.* observed that *Pf*MSP1-specific B cells in adults expressed higher levels of CXCR3 than the total memory B cell population, although the majority of these cells did not have a CD21^-^CD27^-^ phenotype (*48*). In this study, we detected CXCR3^+^ parasite-specific cells but parasite-specific cells were not enriched for CXCR3 expression among atypical B cells. CXCR3 is an interferon-inducible chemokine receptor specific for the chemokines CXCL9, CXCL10, and CXCL11 that are themselves also induced by interferon (*49*). Its function has mainly been studied in T cells where it is required for trafficking of T cells to sites of inflammation (*50*). The profile of trafficking receptor expression in atypical B cells, including CXCR3, has been linked to the propensity of T-bet^hi^ B cells to take up residency in the spleen and bone marrow (*51*). While no data is available for the tissue distribution of atypical B cells in malaria-experienced individuals, it would make sense for parasite-specific B cells to home to the spleen, since this is the main anatomical location for the removal of parasite-infected erythrocytes and cellular debris from the circulation. In this regard, it is interesting to note that the elevated expression of T-bet and CD74 in subset 3 atypical B cells fits the profile of memory B cells that reside in the spleen (*51, 52*), indicating that the loss of these cells in the circulation could be the result of migration into the spleen. However, the observation that the percentage of CXCR3^+^ atypical B cells does not change over time with minimal parasite exposure suggests that the expression of this marker by itself does not direct homing of B cells to tissues. This notion is strengthened by the recent finding that CXCR3 ablation in murine B cells has no effect on the establishment and maintenance of lung-resident memory B cells (*53*). Instead, CXCR3 should probably be considered a marker of activation or prior activation.

One of the complications of studying atypical B cells is that different markers have been used to define atypical B cells. In this study, our definition of atypical B cells is based on how these cells are typically identified in the HIV and malaria fields (CD21^-^CD27^-^) and information from a recent single-cell RNA sequencing study that reported CD11c as the most informative cell surface marker for the identification of atypical B cell lineages (*3*). Our definition of atypical B cells as IgG^+^CD21^-^CD27^-^CD11c^+^ also fits the definition of DN2 B cells, which is how these cells are commonly referred to in the autoimmunity field (IgD^-^CD27^-^CD21^-^ CD11c^+^) and follows recently published recommendations for how these cells should be defined (*22*). Not all of the cells that fall within our definition of atypical B cells are T-bet^+^, which may complicate translating our findings to studies that use T-bet (with or without CD11c) as a defining marker for atypical B cells (*48, 54, 55*).

In conclusion, we found that atypical B cells can be divided into at least three functionally distinct subsets. These findings reconcile many of the conflicting observations that have been made about atypical B cells and provide insight into their function in the immune response against infection. Our study also highlights that the composition of the atypical B cell population can alter with changes in pathogen exposure levels. The selection of individuals and timing of sample collection therefore warrants careful consideration when analyzing atypical B cells and their functional profile. Although the transcriptomes of atypical B cells in chronic infections and autoimmunity are remarkably similar (*11*), our observations about atypical B cells in malaria-experienced individuals will need to be confirmed in other disease models. Collectively, our results suggest that a large fraction of atypical B cells may contribute to a productive and antigen-specific immune response against infection.

## MATERIALS AND METHODS

### Study design and ethics approval

The objective of this study was to assess transcriptional and functional heterogeneity among atypical B cells to determine the function of these cells in the host immune response to infection. Individuals included in this study (n = 29) were residents of the Nagongera sub-county in Tororo District, Uganda. This region was historically characterized by extremely high malaria transmission intensity, with an estimated annual entomological inoculation rate of 125 infectious bites per person per year (*56*). Since 2015, multiple rounds of insecticide spraying (IRS) have dramatically reduced malaria incidence compared with pre-IRS levels (*41*). Individuals were selected for inclusion into this study based on age and *P. falciparum* exposure. In addition, children were selected based on the availability of samples collected at set times relative to a malaria episode (**Table S1**). All individuals included in this study were enrolled in The Program for Resistance, Immunology, Surveillance, and Modeling of Malaria (PRISM) program (*57*) and have provided written consent for the use of their samples for research. The PRISM cohort study was approved by the Makerere University School of Medicine Research and Ethics Committee (SOMREC), London School of Hygiene and Tropical Medicine IRB, the University of California, San Francisco Human Research Protection Program, and the Stanford University School of Medicine IRB. The use of cohort samples for this study was approved by the Institutional Review Board of the University of Texas Health Science Center at San Antonio.

### B cell isolations

Cryopreserved PBMCs were thawed in a 37°C water bath and immediately mixed with pre-warmed thawing medium (IMDM/GlutaMAX supplemented with 10% heat-inactivated FBS (USA origin) and 0.01% Universal Nuclease (Thermo, cat. no. 88700). After centrifugation (250 × g, 5 min), the cell pellet was resuspended in thawing medium. A sample of cells was mixed 1:1 (v/v) with 0.2% trypan blue solution in PBS, and viable cells were counted using a Cellometer Mini (Nexcelom) automated cell counter. Next, cells were centrifuged (250 × g, 5 min) and resuspended in isolation buffer (PBS supplemented with 2% heat-inactivated FBS and 1 mM EDTA) at 50 million live cells/mL and filtered through a 35 µm sterile filter cap (Corning, cat. no. 352235) to break apart aggregated cells. B cells were isolated using the EasySep Human B Cell Isolation Kit (StemCell, cat. no. 17954) or the MojoSort Human Pan B Cell Isolation Kit (BioLegend, cat. no. 480082) according to the manufacturer’s instructions.

### Flow cytometry staining for single-cell analysis

Isolated B cells were washed with PBS (250 × g, 5 min), resuspended in 1 ml of PBS containing 1 µl live/dead stain (LIVE/DEAD Fixable Aqua Dead Cell Stain Kit (Thermo Fisher Scientific, cat. no. L34965), and incubated for 30 min on ice. Subsequently, cells were washed with cold PBS containing 1% bovine serum albumin (BSA) (250 × g, 5 min) and incubated in labeling buffer (total volume of 100 µl) containing PBS, 1% BSA, 5 µl Human TruStain FcX (Biolegend, cat. no. 422301) and 1% dextran sulfate (Sigma-Aldrich, cat. no. 42867-5G) for 10 min at 4°C before incubating a further 30 min at 4°C with an antibody cocktail to label B cell surface markers (**Table S7**). Before acquisition on a BD FACS AriaII cell sorter, cells were washed with cold PBS containing 1% BSA, centrifuged (250 × g, 5 min, 4⁰C), diluted to 20-30 million cells/mL in cold PBS containing 1% BSA, and filtered into a FACS tube through a 35 µm sterile filter cap (Corning, cat. no. 352235) to break apart aggregated cells. Lymphocytes were gated using forward and sideward scatter, followed by doublet exclusion. Cells were then gated on live cells, followed by CD19^+^CD10^-^CD20^+^ B cells. Naive B cells (CD21^+^CD27^-^) and antigen-experienced B cells (all other B cells) were separated based on CD21 and CD27 expression and sorted into cold PBS containing 1% BSA. An average of 37,000 antigen-experienced B cells were mixed with naïve B cells up to a total of 50,000 B cells (**Table S2**). This resulted in an enrichment of antigen-experienced cells from an average of 49% in the original B cell pool to an average of 74% in the final cell suspension. Cells were immediately processed for single-cell sequencing.

### Single-cell gene expression, V(D)J, and feature barcoding library preparation

B cells were counted with a hemocytometer and run on a Chromium 10x controller (10x Genomics) in individual lanes with a targeted recovery rate of 10,000 cells per lane, according to the manufacturer’s instructions. Library preparation was completed following the recommended protocols for Chromium Single Cell 5’ Gene Expression kit as well as 5’ Feature Barcode and V(D)J Enrichment kits (**Table S8**). Libraries were sequenced using the Illumina NovaSeq6000 (paired-end 150 bp) and HiSeq (paired-end 100 bp) platforms.

### Single-cell RNA-seq data analysis

Sequencing reads were mapped to the human reference genome (hg38) and assigned to individual cells of origin using cell-specific barcodes in Cell Ranger (10x Genomics) version 3.1.0. The sequencing data were then processed using Seurat version 3.2.3, following their default pipeline as follows. Low-quality cells with high (>2500) or low (<200) UMI counts were filtered from the samples. In addition, to prevent antibody isotype or V(D)J-gene segment usage from influencing the clustering of cells, we removed all IGH, IGK and IGL transcripts from the single cell data before clustering and UMAP visualization, as proposed by Steward *et al*. (*20*). Gene expression levels for each cell were normalized by total expression, multiplied by a scale factor (10,000), and log-transformed. Cells with > 5% mitochondrial reads were excluded because these are likely damaged or dying cells that have lost much of their cytoplasmic mRNA. Gene expression values for all genes in all clusters are provided in **Table S9**. We determined the top 2000 most variable genes based on their average expression and dispersion. Principal component analysis was performed on highly variable genes to reduce the dimensionality of the data. We performed clustering using the Louvain algorithm implemented in the *FindClusters* function of Seurat (algorithm = 1, resolution = 0.5, dims. use = 1:20) and displayed the clusters using UMAP. One outlier cluster consisting of 13 cells that were most likely T cells was removed from further analysis. Because cells from all samples were distributed evenly across the UMAP (**Figure S3**), no batch corrections were performed. We identified the cluster-specific marker genes conserved across different samples using the *FindConservedMarkers* function (default parameters) from Seurat. To identify differentially expressed genes between a pair of clusters, we used the *FindMarkers* function from Seurat (with parameters *min.pct* = 0.20, and log2FC = 0.25), and additionally filtered genes using a false discovery rate of 0.05. Violin plots, dot plots, and UMAP plots overlaid with gene expression level were generated using standard Seurat functions. Reactome pathway analysis was performed using differentially expressed genes that were upregulated in pairwise cluster comparisons. The top 3 - 5 pathways with FDR-adjusted P value < 0.01 were selected for visual representation in **Figure 3C**. The full set of Reactome analysis results and differentially expressed gene lists are provided in **Tables S5** and **S6**. For heatmaps in Figures 3, gene expression values for the four samples were first divided by the average value for all clusters for that donor to account for donor-to-donor variation. The data points were then z-scored across all clusters and samples prior to plotting heatmaps using the color palette ‘Hiroshige’ from the R package MetBrewer.

### Single-cell RNA-sequencing gene signature score analysis

We performed gene signature score analysis as described in (*11*). Briefly, gene expression data from the Seurat object were scaled (z-score across each gene) to remove bias towards highly expressed genes. A cell-specific signature score was computed by first sorting the normalized scaled gene expression values for each cell followed by summing up the indices (ranks) of the signature genes. These signature scores were then projected onto the transcriptomics-based UMAP.

### Single-cell surface marker analysis

From the single-cell RNA sequencing Seurat object, we extracted the antibody capture component (assay = ADT), normalized the antibody matrix using CLR normalization (Seurat method *NormalizeData*), and then scaled the antibody matrix using Seurat routine *ScaleData*. This normalized and scaled antibody matrix was then used to define the following populations of B cells by phenotype: 1) DN1 B cells: CD27^-^IgG^+^CD21^+^CD11c^-^, 2) DN2 B cells: CD27^-^ IgG^+^CD21^-^CD11c^+^, 3) DN3 B cells: CD27^-^IgG^+^CD21^-^CD11c^-^, 4) DN4 B cells: CD27^-^IgG^+^CD21^+^CD11c^+^, 5) Resting naïve B cells: CD27^-^IgM^+^CD21^+^, 6) Activated naïve B cells: CD27^-^IgM^+^CD21^-^, 7) Resting switched memory B cells: CD27^+^IgG^+^CD21^+^, 8) Activated switched memory B cells: CD27^+^IgG^+^CD21^-^, 9) Resting unswitched memory B cells: CD27^+^IgM^+^CD21^+^, and 10) Activated unswitched memory B cells: CD27^+^IgM^+^CD21^-^. The UMAP plots highlighting corresponding cell categories were generated using Seurat. Distributions of cell surface expression of the markers CD21 and CD27 (**Figure S4**), as well as CD11c and CXCR3 (**Figure S6**) for individual clusters were plotted using the above mentioned normalized and scaled antibody matrix, and employing the 2D kernel density function *kde2d* from the R package *MASS*.

### Single-cell B cell receptor analysis

Single-cell B cell receptor sequencing data was first processed using the command *changeo-10x*. Single-cell heavy chain variable region sequences were then analyzed using the Immcantation pipeline. First, the Immcantation package Change-O (v0.4.4) was used to align lymphocyte receptor sequences to germline sequences for downstream analyses (*58*). Isotype-specific data files were converted into standardized tab-delimited database files required for subsequent Change-O modules to operate using IgBLAST (v1.14.0), which is included in Change-O. The frequency of somatic hypermutations was calculated as described previously (*23*). Average frequencies for each donor and cluster are reported.

To calculate the percentage of clonal expansion within clusters or clonal connections between clusters, genotyping was performed using the routine *tigger-genotype*, followed by analysis of clonally related B cell receptor sequences using a custom Perl script available for download at https://github.com/embunnik/clonal_scores, as described in detail elsewhere (*38*).

### RNA velocity analysis

RNA-velocity analysis was performed by scVelo (*59*), using the scRNA-seq data. First, gene expression matrices from cell ranger were converted to scVelo compatible loom format using the tool Velocyto (*60*). We exported the cluster information and the UMAP coordinates from the Seurat object in .h5ad format, instead of re-computing them in scVelo. RNA-velocity analysis was then performed by the scVelo routine *tl.velocity()* using its default stochastic model. RNA velocity and pseudo-time plots were generated by the default scVelo routines.

### Atypical B cell sorting

B cells were isolated by negative selection as described above, washed with PBS (250 × g, 5 min), and resuspended in 1 ml of PBS containing 1 µl live/dead stain (LIVE/DEAD Fixable Aqua Dead Cell Stain Kit (Thermo Fisher Scientific, cat. no. L34965) and incubated for 30 min on ice. Cells were subsequently washed with cold PBS containing 1% BSA (250 × g, 5 min, 4⁰C) and incubated at 4°C for 30 min with a B cell surface marker antibody cocktail (**Table S12**), added up to a total volume of 100 µl with PBS containing 1% BSA. Cells were then washed with cold PBS containing 1% BSA, centrifuged (250 × g, 5 min, 4⁰C), diluted to 20-30 million cells/mL in cold PBS containing 1% BSA, and filtered into a FACS tube through a 35 µm sterile filter cap (Corning, cat. no. 352235) to break apart aggregated cells. Using a BD FACS Aria Fusion cell sorter, atypical B cell subsets and switched memory cells were sorted into 1.5 mL Eppendorf tubes containing IMDM/GlutaMAX supplemented with 10% heat-inactivated FBS (USA origin). Cells were immediately frozen for RNA-seq or plated for in vitro B cell activation experiments.

### Bulk RNA-Sequencing

RNA-seq libraries were generated following the NEBNext Single Cell/Low Input RNA Library Prep Kit for Illumina (NEB #E6420S) protocol for cells, except that all DNA cleanup steps we done using SparQ magnetic beads (Quantabio). In brief, 200 cells of each B cell subset were FACS sorted directly into 1× NEBNext Cell Lysis Buffer in 1.7 ml low binding tubes (Corning #3207). Cells were then flash frozen in liquid nitrogen for future use. For all samples, cDNA was synthesized and amplified using 21 cycles. cDNA quantity and quality were assessed by Invitrogen Qubit and Agilent 2100 Bioanalyzer. PCR enrichment of adaptor-ligated DNA was performed using NEBext Multiplex Oligos for Ilumina (NEB #E7335S and #E7500S) following the recommended number of amplification cycles based on the amount of cDNA input used in the fragmentation/end prep reaction. Library quantity and quality were assessed by Invitrogen Qubit and Agilent 2100 Bioanalyzer. Eighteen libraries were multiplexed and sequenced on a NovaSeq600 SP 50PE flow cell. Six libraries were retrospectively removed from data analysis due to low input cDNA and resulting low gene expression quality. The list of genes that were differentially expressed between the three atypical B cell clusters in the scRNA-seq data set and were representative of functional differences between these clusters was filtered for an average expression level of 100 transcripts per million (tpm) among the remaining samples. Expression values were z-scored across all samples and plotted in a heatmap.

### Staining for spectral flow analysis

C-terminally biotinylated full-length *P. falciparum* 3D7 MSP1 and AMA1 were produced as described previously (*43*). Antigen tetramers were synthesized by incubating protein with fluorophore-conjugated streptavidin overnight at 4°C at a molar ratio of 6:1 with rotation (**Table S10**). B cells were isolated by negative selection as described above, washed with PBS, centrifuged (250 × g, 5 min), resuspended in 1 ml of PBS containing 1 µl live/dead stain (Zombie UV Fixable Viability kit (Biolegend, cat. no. 423107)) and incubated on ice for 30 min. Cells were subsequently washed with cold PBS containing 1% BSA (250 × g, 5 min, 4⁰C), resuspended with a cocktail of 25 µM of each antigen tetramer diluted in PBS containing 1% BSA to a volume of 100 µl, and incubated at 4°C for 30 min. Next, the cells were washed twice with cold PBS containing 1% BSA (250 × g, 5 min, 4⁰C) and incubated at 4°C for 30 min with a B cell surface marker antibody cocktail (**Table S11**) with 10 µl Brilliant Stain Buffer Plus (BD, cat. No. 566385) diluted in PBS containing 1% BSA up to a volume of 100 µl. The cells were then washed with cold PBS containing 1% BSA (250 × g, 5 min, 4⁰C), resuspended in 1 ml of Transcription Factor Fix/Perm Concentrate (Tonbo, part of cat. no. TNB-0607-KIT), diluted with 3 parts Transcription Factor Fix/Perm Diluent (Tonbo), and incubated at 4°C for 1 hour. After the incubation, the cells were washed twice with 3 ml of 1× Flow Cytometry Perm Buffer (Tonbo) (300 × g, 8 min, 4⁰C) and resuspended in 1× Flow Cytometry Perm Buffer with an intracellular marker antibody (**Table S11**). After an incubation at 4°C for 30 min, the cells were washed twice with 3 ml cold 1× Flow Cytometry Perm Buffer (300 × g, 8 min, 4⁰C) and once with 3 ml cold PBS containing 1% BSA, resuspended in cold PBS containing 1% BSA to 20 – 30 million cells/ml and filtered into a FACS tube through a 35 μm sterile filter cap. Cells were analyzed by flow cytometry immediately following intracellular staining.

### Spectral flow cytometry analysis

B cells were analyzed on a Cytek Aurora spectral flow cytometer equipped with five lasers. SpectroFlo QC Beads (Cytek, cat. no. SKU N7-97355) were run prior to each experiment for performance tracking. Quality control and LJ tracking reports were used to ensure machine performance and that settings between different runs were comparable. Paired samples were processed in the same experiment and experiments performed on separate days contained technical replicates. B cells isolated from pooled PBMCs from two malaria-naïve US donors were used for the compensation of the live/dead stain, for the unstained control, and for technical replicates. Between runs, the expression of cell surface and intracellular markers showed a high correlation between these technical replicates (Spearman r = 0.98, **Figure S19**). UltraComp eBeads Plus Compensation Beads (Thermo, cat. No. 01-3333-41) were used for compensation of all other fluorophores. The cytometry analysis software OMIQ (Dotmatics) was used for the integration and dimension reduction analysis. In short, atypical B cells were first pre-gated on single/live/CD19^+^/CD20^+^/CD21^-^/CD27^-^/CD11c^+^/IgG^+^ cells. UMAPs were then created using the expression of CD19, CD20, CD24, CD38, CD83, CD86, CD95, CXCR3, CXCR5, CD11c, FcRL5, and T-bet as features with default parameters (neighbors = 15, minimum distance = 0.4, metric = Euclidean, random seed = 9258) and included all 38 Ugandan donor samples used in this study for initial projection. To define the three atypical B cell subsets, FlowSOM (*61*) was used to identify three clusters based on CD11c, FcRL5, and CD86 expression using the default parameters (metric = Euclidean, random seed = 1226). For the projection of antigen-specific B cells onto the UMAP, gates were manually set to identify populations of interest using two-dimensional displays, which were then overlaid onto the UMAP projection. Mean fluorescence intensities of cell surface and intracellular markers in select B cell subsets were calculated for each sample individually in OMIQ, and then averaged across samples. To visualize mean fluorescence intensities in a heatmap, the average value for each B cell subset was expressed as a percentage of the highest mean fluorescence intensity among all subsets.

### B cell cultures

One day prior to sorting B cells, wells of a 96-well plate were each seeded with 30,000 adherent, CD40L-expressing 3T3 cells (kind gift from Dr. Mark Connors, NIH) in 100 µl IMDM/Glutamax/ 10% FBS containing 2× MycoZap Plus-PR (Lonza #VZA-2021), 100 ng/ml human IL-2 (GoldBio #1110-02-50), and 100 ng/ml human IL-21 (GoldBio #1110-21-10) to promote expansion and differentiation of B cells into antibody-secreting cells. As a control, one plate was prepared with 100 µl IMDM/Glutamax/ 10% FBS containing 2× MycoZap Plus-PR (Lonza #VZA-2021), 100 ng/ml human IL-2 (GoldBio #1110-02-50), and 100 ng/ml human IL-21 (GoldBio #1110-21-10), without 3T3 cells. Plates were incubated O/N at 37°C and 8% CO_2_. Immediately after B cell sorting, 100 cells of each B cell subset were resuspended in 100 µl IMDM/Glutamax/ 10% FBS, added to the 100 µl culture media with supplements already present in the plates to a total volume of 200 µl, and incubated at 37°C and 8% CO_2_. After seven days, 80 µl of supernatant was removed and replenished with 100 µl IMDM/Glutamax/ 10% FBS containing 2× MycoZap Plus-PR (Lonza #VZA-2021), 100 ng/ml human IL-2 (GoldBio #1110-02-50), 100 ng/ml human IL-21 (GoldBio #1110-21-10). At day 14, the IgG concentration in the supernatant was determined by enzyme-linked immunosorbent assay (ELISA).

### Enzyme-Linked Immunosorbent Assays

To detect IgG, 96-well ELISA plates (Corning #3361) were coated with goat anti-human IgG (Sigma #I2136) antibody at a concentration of 4 µg/ ml diluted in PBS, at a total volume of 100 µl per well. After a one-hour incubation at 37°C or O/N at 4°C, each well was washed once using slowly running (approximately 900 ml/min.) deionized water. All subsequent washes were performed this way. 150 µl blocking buffer (one-third Non-Animal Protein (NAP)-Blocker (G-Biosciences #786-190P) and two-thirds PBS) was added to each well to prevent non-specific binding. After one hour of incubation at 37°C, the wells were washed three times and 5 µl B cell culture supernatant diluted 1:20 in dilution buffer (1% NAP Blocker in PBS; total volume 100 µl) was added per well. Plates were incubated for two hours at 37°C and washed five times. Then, 100 µl 1:2500 diluted (1% NAP Blocker in PBS) HRP-conjugated anti-human IgG antibody (BioLegend #410902) was added to each well. After incubation for one hour at 37°C and three washes, HRP activity was detected using 50 µl TMB (Thermo #PI34024). Plates were incubated in the dark at RT and the oxidation reaction was stopped by adding 50 µl 0.18 M H_2_SO_4_ (Fisher #FLA300-212) per well when the negative controls (wells that received buffer when test wells received culture supernatant) started to color. Absorbance was measured at 450 nm using a BioTek Synergy H4 microplate reader. A human IgG (Sigma #I2511) standard curve (ten three-fold serial dilutions starting at 20 µg/ml) was used to quantify samples.

## Acknowledgements

This work was supported by the National Institutes of Health (R01 AI153425 to EMB, R35 GM128938 to FA, F31 AI169993 to RAR, TL1 TR002647 to RAR, T32 AI138944 to ABR, U19 AI150741 to BG and PJ, and U19 AI089674). The authors acknowledge the Cell Analysis Core Facility (Dr. Sandra Cardona) and the Genomics Research Core Facility (Sean Vargas) at the University of Texas at San Antonio for support during this work. Data were generated in the Genome Sequencing Facility, which is supported by UT Health San Antonio, NIH-NCI P30 CA054174 (Mays Cancer Center at UT Health San Antonio), and NIH Shared Instrument grant 1S10OD030311-01 (S10 grant towards NovaSeq6000 sequencing system) and in the Flow Cytometry Shared Resource Facility, which is supported by UT Health San Antonio, NIH-NCI P30 CA054174 (Mays Cancer Center at UT Health), CPRIT (RP210126) and NIH Shared Instrument grant 1S10OD030432-01 (S10 grant towards the purchase of a DB FACSAria Fusion). Plasmids encoding 3D7 MSP1-bio, AMA1-bio, and BirA, were a kind gift from Dr. Gavin Wright (Wellcome Sanger Institute; Addgene plasmids #47709, 47741, and 32408).

## Author contributions

GB performed single-cell sequencing experiments and data analysis. RAR performed bulk RNA-sequencing experiments, spectral flow cytometry experiments, *in vitro* B cell cultures, ELISAs and data analysis. PD and FA analyzed single-cell sequencing data. ABR analyzed somatic hypermutation data. RG provided critical feedback on the manuscript. IS, PJ, MEF, and BG provided clinical samples. RAR, GB, SB, and EMB wrote the manuscript with input from all other co-authors. All authors contributed to the article and approved the submitted version.

## Data availability

All sequencing files will be available from the NCBI Sequencing Read Archive under BioProject accession no. TBD.

## References and notes

1. D. Lau, L. Y. Lan, S. F. Andrews, C. Henry, K. T. Rojas, K. E. Neu, M. Huang, Y. Huang, B. DeKosky, A. E. Palm, G. C. Ippolito, G. Georgiou, P. C. Wilson, Low CD21 expression defines a population of recent germinal center graduates primed for plasma cell differentiation. Sci Immunol 2, (2017).

2. S. F. Andrews, M. J. Chambers, C. A. Schramm, J. Plyler, J. E. Raab, M. Kanekiyo, R. Gillespie, A. Ransier, S. Darko, J. Hu, X. Chen, H. M. Yassine, J. C. Boyington, M. C. Crank, G. L. Chen, E. Coates, J. R. Mascola, D. C. Douek, B. S. Graham, J. E. Ledgerwood, A. B. McDermott, Activation Dynamics and Immunoglobulin Evolution of Pre-existing and Newly Generated Human Memory B cell Responses to Influenza Hemagglutinin. Immunity 51, 398–410 e395 (2019).

3. H. J. Sutton, R. Aye, A. H. Idris, R. Vistein, E. Nduati, O. Kai, J. Mwacharo, X. Li, X. Gao, T. D. Andrews, M. Koutsakos, T. H. O. Nguyen, M. Nekrasov, P. Milburn, A. Eltahla, A. A. Berry, N. Kc, S. Chakravarty, B. K. L. Sim, A. K. Wheatley, S. J. Kent, S. L. Hoffman, K. E. Lyke, P. Bejon, F. Luciani, K. Kedzierska, R. A. Seder, F. M. Ndungu, I. Cockburn, Atypical B cells are part of an alternative lineage of B cells that participates in responses to vaccination and infection in humans. Cell Rep 34, 108684 (2021).

4. C. Sundling, C. Ronnberg, V. Yman, M. Asghar, P. Jahnmatz, T. Lakshmikanth, Y. Chen, J. Mikes, M. N. Forsell, K. Sonden, A. Achour, P. Brodin, K. E. Persson, A. Farnert, B cell profiling in malaria reveals expansion and remodelling of CD11c+ B cell subsets. JCI Insight 5, (2019).

5. S. Moir, J. Ho, A. Malaspina, W. Wang, A. C. DiPoto, M. A. O’Shea, G. Roby, S. Kottilil, J. Arthos, M. A. Proschan, T. W. Chun, A. S. Fauci, Evidence for HIV-associated B cell exhaustion in a dysfunctional memory B cell compartment in HIV-infected viremic individuals. J Exp Med 205, 1797–1805 (2008).

6. G. E. Weiss, P. D. Crompton, S. Li, L. A. Walsh, S. Moir, B. Traore, K. Kayentao, A. Ongoiba, O. K. Doumbo, S. K. Pierce, Atypical memory B cells are greatly expanded in individuals living in a malaria-endemic area. J Immunol 183, 2176–2182 (2009).

7. C. Wei, J. Anolik, A. Cappione, B. Zheng, A. Pugh-Bernard, J. Brooks, E. H. Lee, E. C. Milner, I. Sanz, A new population of cells lacking expression of CD27 represents a notable component of the B cell memory compartment in systemic lupus erythematosus. J Immunol 178, 6624–6633 (2007).

8. I. Isnardi, Y. S. Ng, L. Menard, G. Meyers, D. Saadoun, I. Srdanovic, J. Samuels, J. Berman, J. H. Buckner, C. Cunningham-Rundles, E. Meffre, Complement receptor 2/CD21-human naive B cells contain mostly autoreactive unresponsive clones. Blood 115, 5026–5036 (2010).

9. S. A. Jenks, K. S. Cashman, E. Zumaquero, U. M. Marigorta, A. V. Patel, X. Wang, D. Tomar, M. C. Woodruff, Z. Simon, R. Bugrovsky, E. L. Blalock, C. D. Scharer, C. M. Tipton, C. Wei, S. S. Lim, M. Petri, T. B. Niewold, J. H. Anolik, G. Gibson, F. E. Lee, J. M. Boss, F. E. Lund, I. Sanz, Distinct Effector B Cells Induced by Unregulated Toll-like Receptor 7 Contribute to Pathogenic Responses in Systemic Lupus Erythematosus. Immunity 49, 725–739 e726 (2018).

10. R. T. Sullivan, C. C. Kim, M. F. Fontana, M. E. Feeney, P. Jagannathan, M. J. Boyle, C. J. Drakeley, I. Ssewanyana, F. Nankya, H. Mayanja-Kizza, G. Dorsey, B. Greenhouse, FCRL5 Delineates Functionally Impaired Memory B Cells Associated with Plasmodium falciparum Exposure. PLoS Pathog 11, e1004894 (2015).

11. P. Holla, B. Dizon, A. A. Ambegaonkar, N. Rogel, E. Goldschmidt, A. K. Boddapati, H. Sohn, D. Sturdevant, J. W. Austin, L. Kardava, L. Yuesheng, P. Liu, S. Moir, S. K. Pierce, A. Madi, Shared transcriptional profiles of atypical B cells suggest common drivers of expansion and function in malaria, HIV, and autoimmunity. Sci Adv 7, (2021).

12. S. Portugal, C. M. Tipton, H. Sohn, Y. Kone, J. Wang, S. Li, J. Skinner, K. Virtaneva, D. E. Sturdevant, S. F. Porcella, O. K. Doumbo, S. Doumbo, K. Kayentao, A. Ongoiba, B. Traore, I. Sanz, S. K. Pierce, P. D. Crompton, Malaria-associated atypical memory B cells exhibit markedly reduced B cell receptor signaling and effector function. Elife 4, (2015).

13. A. A. Ambegaonkar, K. Kwak, H. Sohn, J. Manzella-Lapeira, J. Brzostowski, S. K. Pierce, Expression of inhibitory receptors by B cells in chronic human infectious diseases restricts responses to membrane-associated antigens. Sci Adv 6, eaba6493 (2020).

14. E. Wing, C. Sutherland, K. Miles, D. Gray, C. Goodyear, T. D. Otto, S. Breusch, G. J. M. Cowan, M. Gray, A comprehensive analysis of rheumatoid arthritis B cells reveals the importance of CD11c^+ve^ double-negative-2 B cells as the major synovial plasma cell precursor. bioRxiv, 2023.2002.2015.526468 (2023).

15. M. F. Muellenbeck, B. Ueberheide, B. Amulic, A. Epp, D. Fenyo, C. E. Busse, M. Esen, M. Theisen, B. Mordmuller, H. Wardemann, Atypical and classical memory B cells produce Plasmodium falciparum neutralizing antibodies. J Exp Med 210, 389–399 (2013).

16. P. Holla, A. Ambegaonkar, H. Sohn, S. K. Pierce, Exhaustion may not be in the human B cell vocabulary, at least not in malaria. Immunol Rev 292, 139–148 (2019).

17. C. S. Hopp, J. Skinner, S. L. Anzick, C. M. Tipton, M. E. Peterson, S. Li, S. Doumbo, K. Kayentao, A. Ongoiba, C. Martens, B. Traore, P. D. Crompton, Atypical B cells up-regulate costimulatory molecules during malaria and secrete antibodies with T follicular helper cell support. Sci Immunol 7, eabn1250 (2022).

18. A. E. Braddom, G. Batugedara, S. Bol, E. M. Bunnik, Potential functions of atypical memory B cells in Plasmodium-exposed individuals. Int J Parasitol 50, 1033–1042 (2020).

19. C. M. Andrade, H. Fleckenstein, R. Thomson-Luque, S. Doumbo, N. F. Lima, C. Anderson, J. Hibbert, C. S. Hopp, T. M. Tran, S. Li, M. Niangaly, H. Cisse, D. Doumtabe, J. Skinner, D. Sturdevant, S. Ricklefs, K. Virtaneva, M. Asghar, M. V. Homann, L. Turner, J. Martins, E. L. Allman, M. E. N’Dri, V. Winkler, M. Llinas, C. Lavazec, C. Martens, A. Farnert, K. Kayentao, A. Ongoiba, T. Lavstsen, N. S. Osorio, T. D. Otto, M. Recker, B. Traore, P. D. Crompton, S. Portugal, Increased circulation time of Plasmodium falciparum underlies persistent asymptomatic infection in the dry season. Nat Med 26, 1929–1940 (2020).

20. A. Stewart, J. C. Ng, G. Wallis, V. Tsioligka, F. Fraternali, D. K. Dunn-Walters, Single-Cell Transcriptomic Analyses Define Distinct Peripheral B Cell Subsets and Discrete Development Pathways. Front Immunol 12, 602539 (2021).

21. A. H. Ellebedy, K. J. Jackson, H. T. Kissick, H. I. Nakaya, C. W. Davis, K. M. Roskin, A. K. McElroy, C. M. Oshansky, R. Elbein, S. Thomas, G. M. Lyon, C. F. Spiropoulou, A. K. Mehta, P. G. Thomas, S. D. Boyd, R. Ahmed, Defining antigen-specific plasmablast and memory B cell subsets in human blood after viral infection or vaccination. Nat Immunol 17, 1226–1234 (2016).

22. I. Sanz, C. Wei, S. A. Jenks, K. S. Cashman, C. Tipton, M. C. Woodruff, J. Hom, F. E. Lee, Challenges and Opportunities for Consistent Classification of Human B Cell and Plasma Cell Populations. Front Immunol 10, 2458 (2019).

23. A. E. Braddom, S. Bol, S. J. Gonzales, R. A. Reyes, K. Musinguzi, F. Nankya, I. Ssewanyana, B. Greenhouse, E. M. Bunnik, B Cell Receptor Repertoire Analysis in Malaria-Naive and Malaria-Experienced Individuals Reveals Unique Characteristics of Atypical Memory B Cells. mSphere 6, e0072621 (2021).

24. D. R. Glass, A. G. Tsai, J. P. Oliveria, F. J. Hartmann, S. C. Kimmey, A. A. Calderon, L. Borges, M. C. Glass, L. E. Wagar, M. M. Davis, S. C. Bendall, An Integrated Multi-omic Single-Cell Atlas of Human B Cell Identity. Immunity 53, 217–232 e215 (2020).

25. S. Gu, X. Han, C. Yao, H. Ding, J. Zhang, G. Hou, B. Qu, H. Zhou, Z. Ying, Z. Ye, J. Qian, Q. Guo, S. Chen, C. G. Vinuesa, D. Dai, N. Shen, The transcription factor Zeb2 drives differentiation of age-associated B cells. bioRxiv, 2021.2007.2024.453633 (2021).

26. X. Gao, Q. Shen, J. A. Roco, K. Frith, C. M. L. Munier, M. Nekrasov, B. Dalton, J.-S. He, R. Jaeger, M. C. Cook, J. J. Zaunders, I. A. Cockburn, ZEB2 regulates the development of CD11c+ atypical B cells. bioRxiv, 2022.2009.2001.506173 (2022).

27. C. Akatsu, K. Shinagawa, N. Numoto, Z. Liu, A. K. Ucar, M. Aslam, S. Phoon, T. Adachi, K. Furukawa, N. Ito, T. Tsubata, CD72 negatively regulates B lymphocyte responses to the lupus-related endogenous toll-like receptor 7 ligand Sm/RNP. J Exp Med 213, 2691–2706 (2016).

28. C. Tan, R. Hiwa, J. L. Mueller, V. Vykunta, K. Hibiya, M. Noviski, J. Huizar, J. F. Brooks, J. Garcia, C. Heyn, Z. Li, A. Marson, J. Zikherman, NR4A nuclear receptors restrain B cell responses to antigen when second signals are absent or limiting. Nat Immunol 21, 1267–1279 (2020).

29. C. Tan, J. L. Mueller, M. Noviski, J. Huizar, D. Lau, A. Dubinin, A. Molofsky, P. C. Wilson, J. Zikherman, Nur77 Links Chronic Antigen Stimulation to B Cell Tolerance by Restricting the Survival of Self-Reactive B Cells in the Periphery. J Immunol 202, 2907–2923 (2019).

30. J. Campisi, Aging, cellular senescence, and cancer. Annu Rev Physiol 75, 685–705 (2013).

31. K. Contrepois, C. Coudereau, B. A. Benayoun, N. Schuler, P. F. Roux, O. Bischof, R. Courbeyrette, C. Carvalho, J. Y. Thuret, Z. Ma, C. Derbois, M. C. Nevers, H. Volland, C. E. Redon, W. M. Bonner, J. F. Deleuze, C. Wiel, D. Bernard, M. P. Snyder, C. E. Rube, R. Olaso, F. Fenaille, C. Mann, Histone variant H2A.J accumulates in senescent cells and promotes inflammatory gene expression. Nat Commun 8, 14995 (2017).

32. D. Frasca, M. Romero, A. Diaz, D. Garcia, S. Thaller, B. B. Blomberg, B Cells with a Senescent-Associated Secretory Phenotype Accumulate in the Adipose Tissue of Individuals with Obesity. Int J Mol Sci 22, (2021).

33. M. G. Giudizi, R. Biagiotti, F. Almerigogna, A. Alessi, A. Tiri, G. F. Del Prete, S. Ferrone, S. Romagnani, Role of HLA class I and class II antigens in activation and differentiation of B cells. Cell Immunol 108, 97–108 (1987).

34. J. G. Cyster, C. D. C. Allen, B Cell Responses: Cell Interaction Dynamics and Decisions. Cell 177, 524–540 (2019).

35. P. Stumptner-Cuvelette, P. Benaroch, Multiple roles of the invariant chain in MHC class II function. Biochim Biophys Acta 1542, 1–13 (2002).

36. R. Di Niro, S. J. Lee, J. A. Vander Heiden, R. A. Elsner, N. Trivedi, J. M. Bannock, N. T. Gupta, S. H. Kleinstein, F. Vigneault, T. J. Gilbert, E. Meffre, S. J. McSorley, M. J. Shlomchik, Salmonella Infection Drives Promiscuous B Cell Activation Followed by Extrafollicular Affinity Maturation. Immunity 43, 120–131 (2015).

37. V. Bergen, M. Lange, S. Peidli, F. A. Wolf, F. J. Theis, Generalizing RNA velocity to transient cell states through dynamical modeling. Nat Biotechnol 38, 1408–1414 (2020).

38. S. J. Gonzales, S. Bol, A. E. Braddom, R. Sullivan, R. A. Reyes, I. Ssewanyana, E. Eggers, B. Greenhouse, E. M. Bunnik, Longitudinal analysis of FcRL5 expression and clonal relationships among classical and atypical memory B cells following malaria. Malar J 20, 435 (2021).

39. U. Hershberg, E. T. Luning Prak, The analysis of clonal expansions in normal and autoimmune B cell repertoires. Philos Trans R Soc Lond B Biol Sci 370, (2015).

40. W. Meng, B. Zhang, G. W. Schwartz, A. M. Rosenfeld, D. Ren, J. J. C. Thome, D. J. Carpenter, N. Matsuoka, H. Lerner, A. L. Friedman, T. Granot, D. L. Farber, M. J. Shlomchik, U. Hershberg, E. T. Luning Prak, An atlas of B-cell clonal distribution in the human body. Nat Biotechnol 35, 879–884 (2017).

41. K. Zinszer, K. Charland, S. Vahey, D. Jahagirdar, J. C. Rek, E. Arinaitwe, J. Nankabirwa, K. Morrison, M. L. Sadoine, M. A. Tutt-Guerette, S. G. Staedke, M. R. Kamya, B. Greenhouse, I. Rodriguez-Barraquer, G. Dorsey, The Impact of Multiple Rounds of Indoor Residual Spraying on Malaria Incidence and Hemoglobin Levels in a High-Transmission Setting. J Infect Dis 221, 304–312 (2020).

42. M. R. Kamya, A. Kakuru, M. Muhindo, E. Arinaitwe, J. I. Nankabirwa, J. Rek, V. Bigira, J. Kapisi, H. Wanzira, J. Achan, P. Natureeba, A. Gasasira, D. Havlir, P. Jagannathan, P. J. Rosenthal, I. Rodriguez-Barraquer, G. Dorsey, The Impact of Control Interventions on Malaria Burden in Young Children in a Historically High-Transmission District of Uganda: A Pooled Analysis of Cohort Studies from 2007 to 2018. Am J Trop Med Hyg 103, 785–792 (2020).

43. S. J. Gonzales, K. N. Clarke, G. Batugedara, R. Garza, A. E. Braddom, R. A. Reyes, I. Ssewanyana, K. C. Garrison, G. C. Ippolito, B. Greenhouse, S. Bol, E. M. Bunnik, A Molecular Analysis of Memory B Cell and Antibody Responses Against Plasmodium falciparum Merozoite Surface Protein 1 in Children and Adults From Uganda. Front Immunol 13, 809264 (2022).

44. R. A. Reyes, K. Clarke, S. J. Gonzales, A. M. Cantwell, R. Garza, G. Catano, R. E. Tragus, T. F. Patterson, S. Bol, E. M. Bunnik, SARS-CoV-2 spike-specific memory B cells express higher levels of T-bet and FcRL5 after non-severe COVID-19 as compared to severe disease. PLoS One 16, e0261656 (2021).

45. C. L. Holness, D. L. Simmons, Molecular cloning of CD68, a human macrophage marker related to lysosomal glycoproteins. Blood 81, 1607–1613 (1993).

46. E. Gottfried, L. A. Kunz-Schughart, A. Weber, M. Rehli, A. Peuker, A. Muller, M. Kastenberger, G. Brockhoff, R. Andreesen, M. Kreutz, Expression of CD68 in non-myeloid cell types. Scand J Immunol 67, 453–463 (2008).

47. C. S. Hopp, J. Skinner, S. L. Anzick, C. M. Tipton, M. E. Peterson, L. Shanping, S. Doumbo, K. Kayentao, A. Ongoiba, C. Martens, B. Traore, P. D. Crompton, Atypical B cells upregulate co-stimulatory molecules during malaria and secrete antibodies with T follicular helper cell support. BioRxiv, (2021).

48. J. L. Johnson, R. L. Rosenthal, J. J. Knox, A. Myles, M. S. Naradikian, J. Madej, M. Kostiv, A. M. Rosenfeld, W. Meng, S. R. Christensen, S. E. Hensley, J. Yewdell, D. H. Canaday, J. Zhu, A. B. McDermott, Y. Dori, M. Itkin, E. J. Wherry, N. Pardi, D. Weissman, A. Naji, E. T. L. Prak, M. R. Betts, M. P. Cancro, The Transcription Factor T-bet Resolves Memory B Cell Subsets with Distinct Tissue Distributions and Antibody Specificities in Mice and Humans. Immunity 52, 842–855 e846 (2020).

49. J. R. Groom, A. D. Luster, CXCR3 ligands: redundant, collaborative and antagonistic functions. Immunol Cell Biol 89, 207–215 (2011).

50. J. R. Groom, A. D. Luster, CXCR3 in T cell function. Exp Cell Res 317, 620–631 (2011).

51. G. Fontanesi, P. Costa, R. Rotini, P. Pignedoli, Total P.C.A. knee arthroprosthesis. Ital J Orthop Traumatol 14, 41–48 (1988).

52. N. M. Weisel, S. M. Joachim, S. Smita, D. Callahan, R. A. Elsner, L. J. Conter, M. Chikina, D. L. Farber, F. J. Weisel, M. J. Shlomchik, Surface phenotypes of naive and memory B cells in mouse and human tissues. Nat Immunol 23, 135–145 (2022).

53. C. Gregoire, L. Spinelli, S. Villazala-Merino, L. Gil, M. P. Holgado, M. Moussa, C. Dong, A. Zarubica, M. Fallet, J. M. Navarro, B. Malissen, P. Milpied, M. Gaya, Viral infection engenders bona fide and bystander subsets of lung-resident memory B cells through a permissive mechanism. Immunity 55, 1216–1233 e1219 (2022).

54. B. Keller, V. Strohmeier, I. Harder, S. Unger, K. J. Payne, G. Andrieux, M. Boerries, P. T. Felixberger, J. J. M. Landry, A. Nieters, A. Rensing-Ehl, U. Salzer, N. Frede, S. Usadel, R. Elling, C. Speckmann, I. Hainmann, E. Ralph, K. Gilmour, M. W. J. Wentink, M. van der Burg, H. S. Kuehn, S. D. Rosenzweig, U. Kolsch, H. von Bernuth, P. Kaiser-Labusch, F. Gothe, S. Hambleton, A. D. Vlagea, A. Garcia Garcia, L. Alsina, G. Markelj, T. Avcin, J. Vasconcelos, M. Guedes, J. Y. Ding, C. L. Ku, B. Shadur, D. T. Avery, N. Venhoff, J. Thiel, H. Becker, L. Erazo-Borras, C. M. Trujillo-Vargas, J. L. Franco, C. Fieschi, S. Okada, P. E. Gray, G. Uzel, J. L. Casanova, M. Fliegauf, B. Grimbacher, H. Eibel, S. Ehl, R. E. Voll, M. Rizzi, P. Stepensky, V. Benes, C. S. Ma, C. Bossen, S. G. Tangye, K. Warnatz, The expansion of human T-bet(high)CD21(low) B cells is T cell dependent. Sci Immunol 6, eabh0891 (2021).

55. S. Wang, J. Wang, V. Kumar, J. L. Karnell, B. Naiman, P. S. Gross, S. Rahman, K. Zerrouki, R. Hanna, C. Morehouse, N. Holoweckyj, H. Liu, T. Autoimmunity Molecular Medicine, Z. Manna, R. Goldbach-Mansky, S. Hasni, R. Siegel, M. Sanjuan, K. Streicher, M. P. Cancro, R. Kolbeck, R. Ettinger, IL-21 drives expansion and plasma cell differentiation of autoreactive CD11c(hi)T-bet(+) B cells in SLE. Nat Commun 9, 1758 (2018).

56. M. Kilama, D. L. Smith, R. Hutchinson, R. Kigozi, A. Yeka, G. Lavoy, M. R. Kamya, S. G. Staedke, M. J. Donnelly, C. Drakeley, B. Greenhouse, G. Dorsey, S. W. Lindsay, Estimating the annual entomological inoculation rate for Plasmodium falciparum transmitted by Anopheles gambiae s.l. using three sampling methods in three sites in Uganda. Malar J 13, 111 (2014).

57. M. R. Kamya, E. Arinaitwe, H. Wanzira, A. Katureebe, C. Barusya, S. P. Kigozi, M. Kilama, A. J. Tatem, P. J. Rosenthal, C. Drakeley, S. W. Lindsay, S. G. Staedke, D. L. Smith, B. Greenhouse, G. Dorsey, Malaria transmission, infection, and disease at three sites with varied transmission intensity in Uganda: implications for malaria control. Am J Trop Med Hyg 92, 903–912 (2015).

58. N. T. Gupta, J. A. Vander Heiden, M. Uduman, D. Gadala-Maria, G. Yaari, S. H. Kleinstein, Change-O: a toolkit for analyzing large-scale B cell immunoglobulin repertoire sequencing data. Bioinformatics 31, 3356–3358 (2015).

59. E. Chichin, Community care for the frail elderly: the case of non-professional home care workers. Women Health 14, 93–104 (1988).

60. G. La Manno, R. Soldatov, A. Zeisel, E. Braun, H. Hochgerner, V. Petukhov, K. Lidschreiber, M. E. Kastriti, P. Lonnerberg, A. Furlan, J. Fan, L. E. Borm, Z. Liu, D. van Bruggen, J. Guo, X. He, R. Barker, E. Sundstrom, G. Castelo-Branco, P. Cramer, I. Adameyko, S. Linnarsson, P. V. Kharchenko, RNA velocity of single cells. Nature 560, 494–498 (2018).

61. S. Van Gassen, B. Callebaut, M. J. Van Helden, B. N. Lambrecht, P. Demeester, T. Dhaene, Y. Saeys, FlowSOM: Using self-organizing maps for visualization and interpretation of cytometry data. Cytometry A 87, 636–645 (2015).

