## Supplementary Figures and Tables for "Atypical B cells consist of subsets with distinct effector functions"

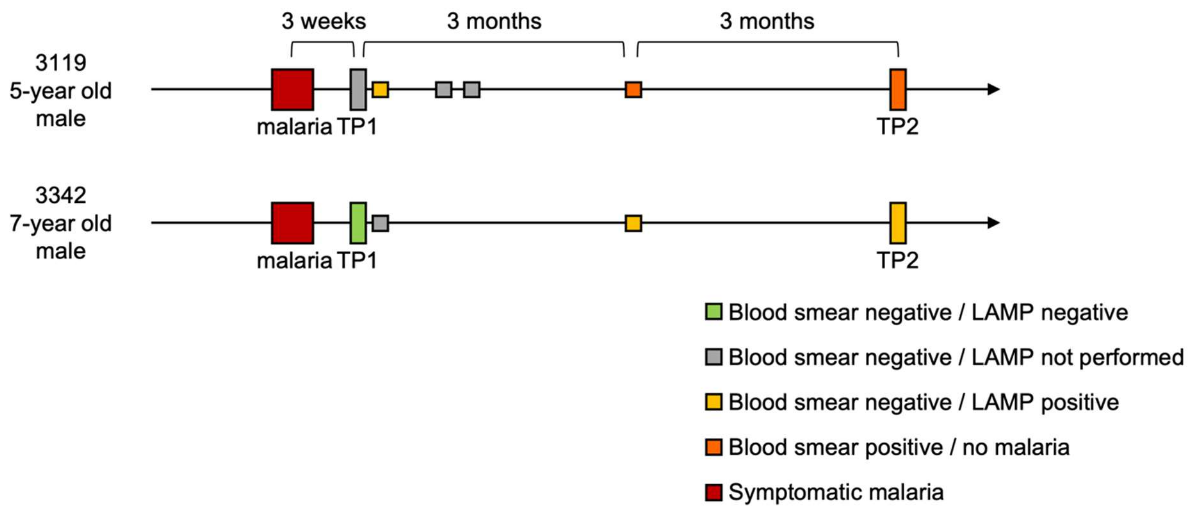


**Figure S1: Schematic overview of *P. falciparum* infection history.** The outcome of screening for parasitemia between the two sampling time points is shown. TP, time point; LAMP, loop-mediated isothermal amplification assay.

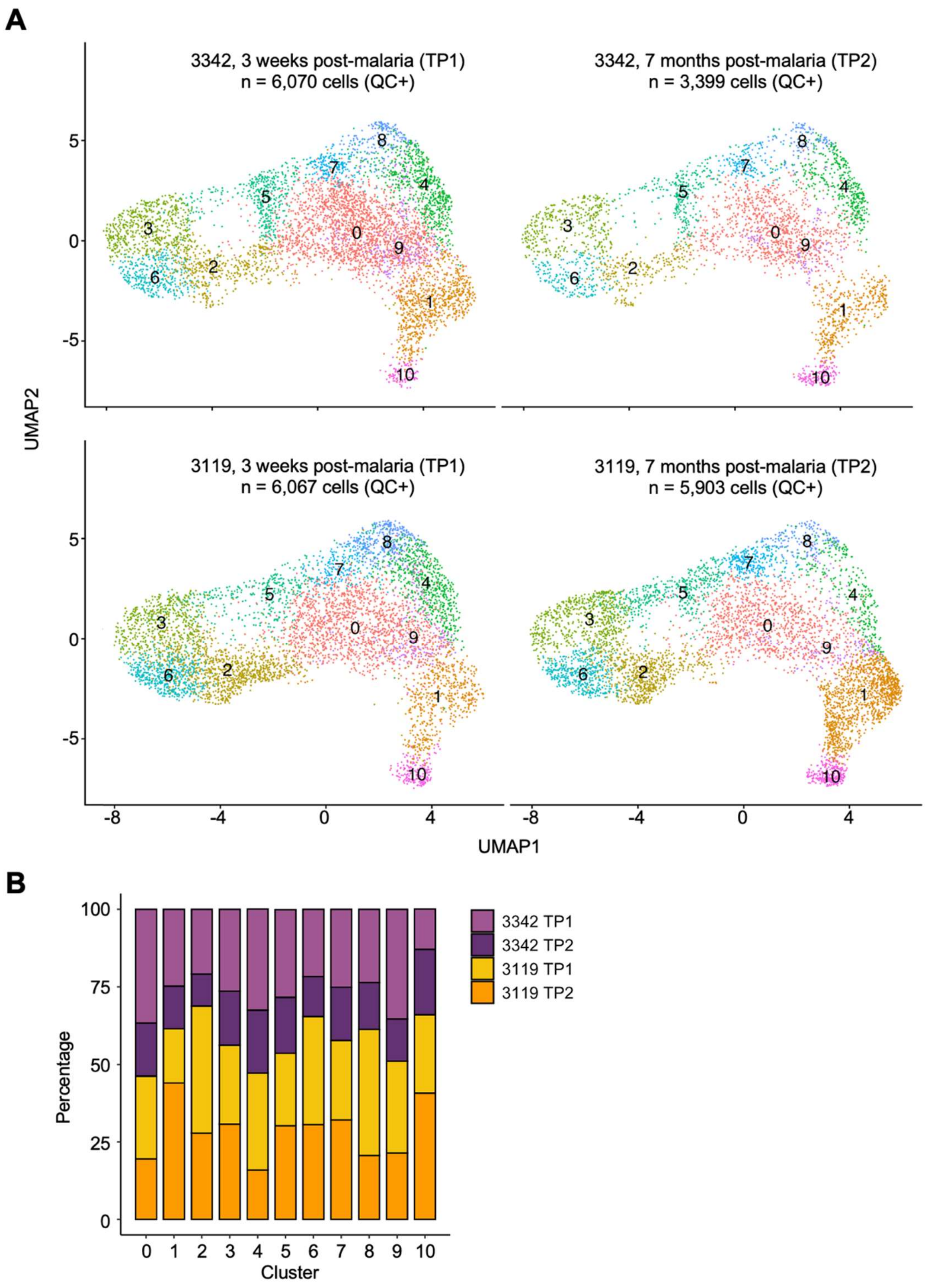


**Figure S2: Contribution of individual samples to the total data set. A)** B cells that passed quality control (QC+) from each individual sample shown in the composite UMAP. **B)** The percentage of cells from each sample in each cluster. See table S4 for absolute numbers.

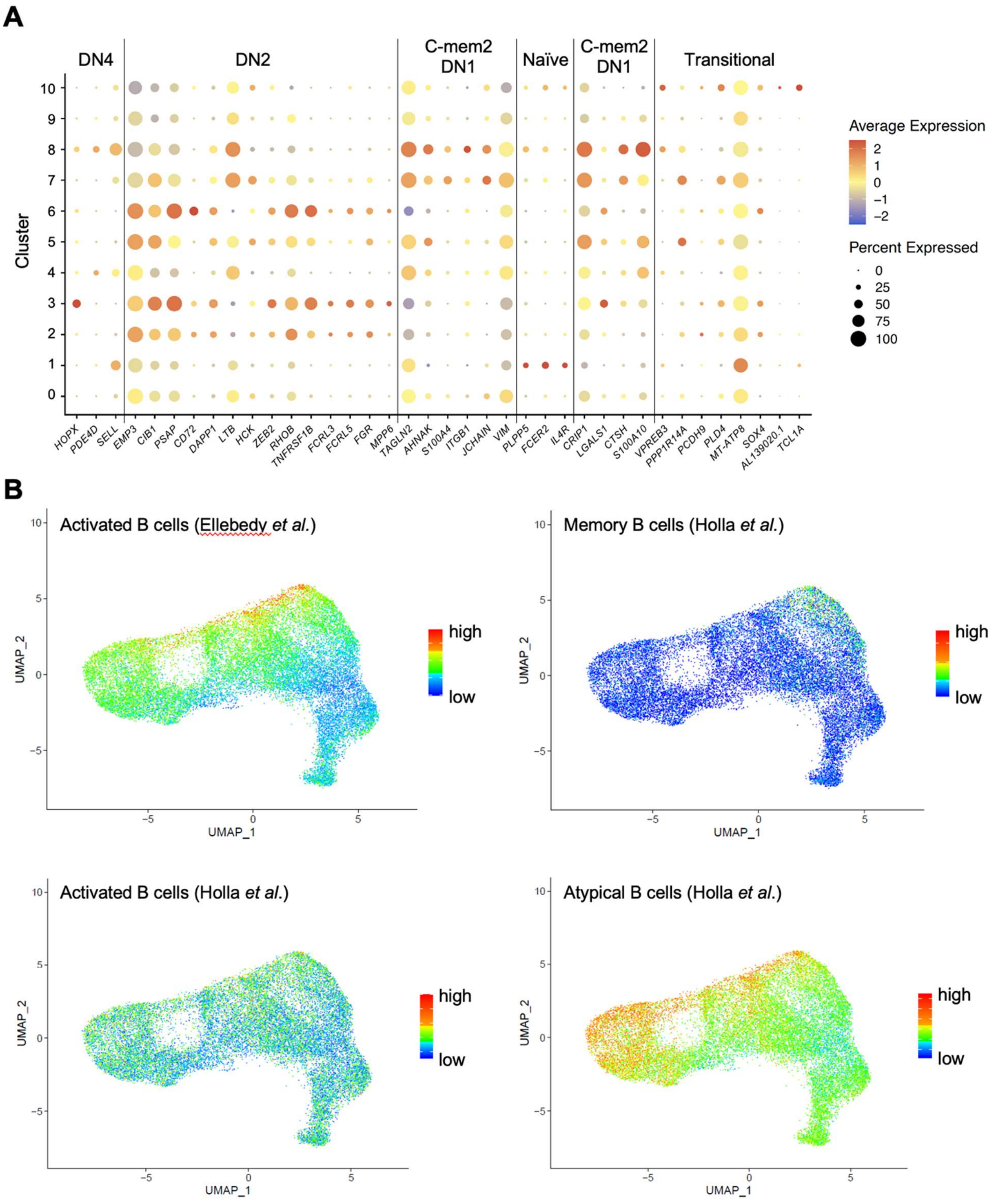
A
DN4
DN2
C-mem2 DN1
Naïve
C-mem2 DN1
Transitional
Cluster
Average Expression
Percent Expressed
HOPX
PDE4D
SELL
EMP3
CIB1
PSAP
CD72
DAPP1
LTB
HCK
ZEB2
RHOB
TNFRSF1B
FCRL3
FCRL5
FGR
MPP6
TAGLN2
AHNAK
S100A4
ITGB1
JCHAIN
VIM
PLPP5
FCER2
IL4R
CRIP1
LGALS1
CTSH
S100A10
VPREB3
PPP1R14A
PCDH9
PLD4
MT-ATP8
SOX4
AL139020.1
TCL1A
B
Activated B cells (Ellebedy et al.)
Memory B cells (Holla et al.)
Activated B cells (Holla et al.)
Atypical B cells (Holla et al.)
UMAP_1
UMAP_2
high
low

**Figure S3: Comparisons between B cell clusters identified in this study and previously published gene signatures. A)** For each cluster, average expression levels are shown for genes that were most highly differentially expressed in B cell subsets identified by Stewart et al. (compare to Fig. 1C in (*20*)). The corresponding subsets from Stewart et al. are indicated in the top. B) Gene signatures from $CD71^+$ activated B cells (Ellebedy et al. (*21*)) and memory B cells, activated B cells, and atypical B cells (all from Holla et al. (*11*)) projected onto the UMAP. Cell-specific signature scores were calculated by first ranking the normalized z-scored gene expression values for each cell, followed by adding up the ranks of the signature genes, as described in (*11*). Activated B cells from Holla et al. show a uniform pattern across the UMAP. We therefore used the activated B cell signature from Ellebedy et al. to broadly define the location of these cells.

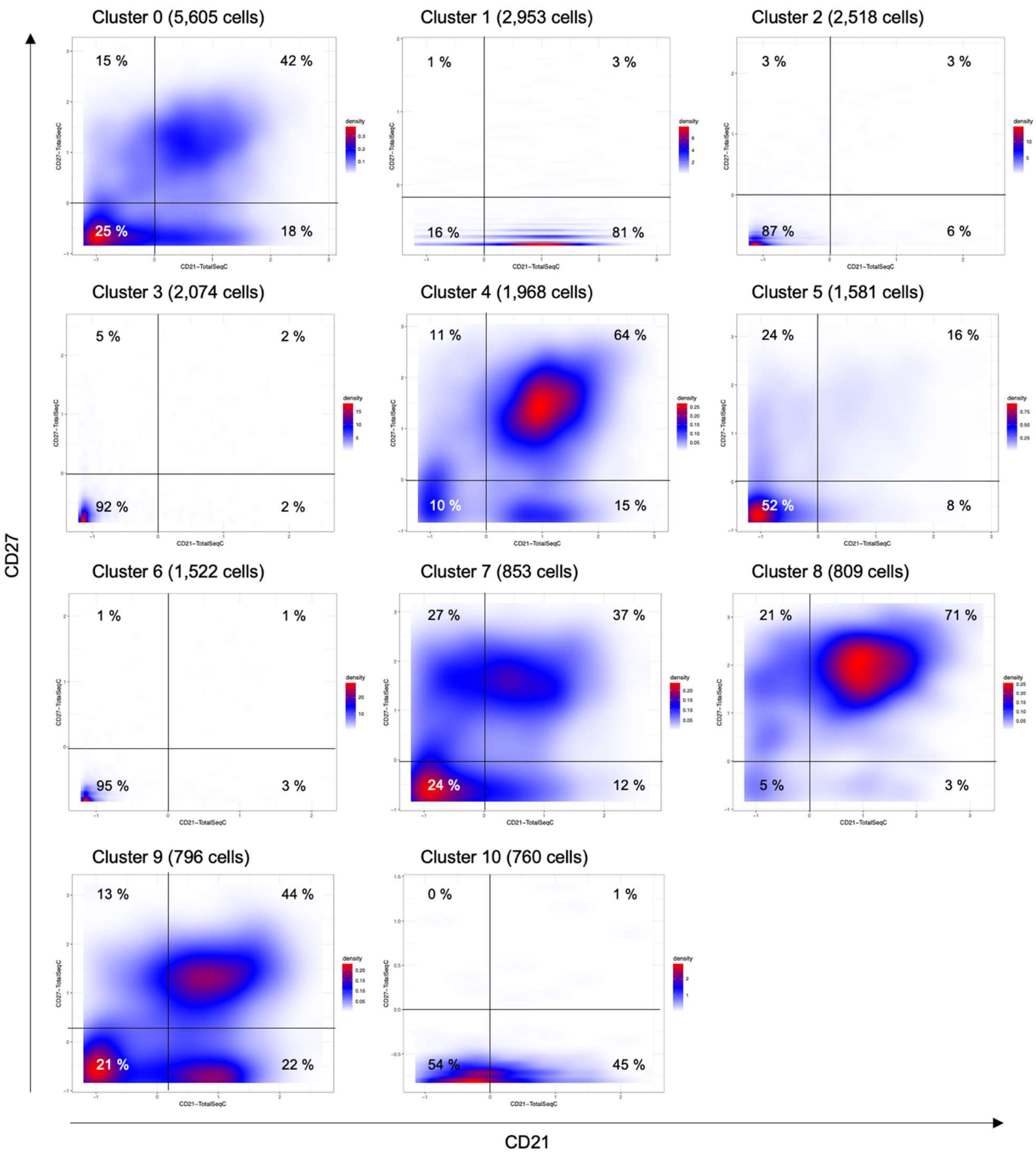


**Figure S4: Cell surface expression of CD21 and CD27 in each cluster as determined by feature barcoding.** Lines in the graphs denote the cutoffs for expression (normalized values > 0).

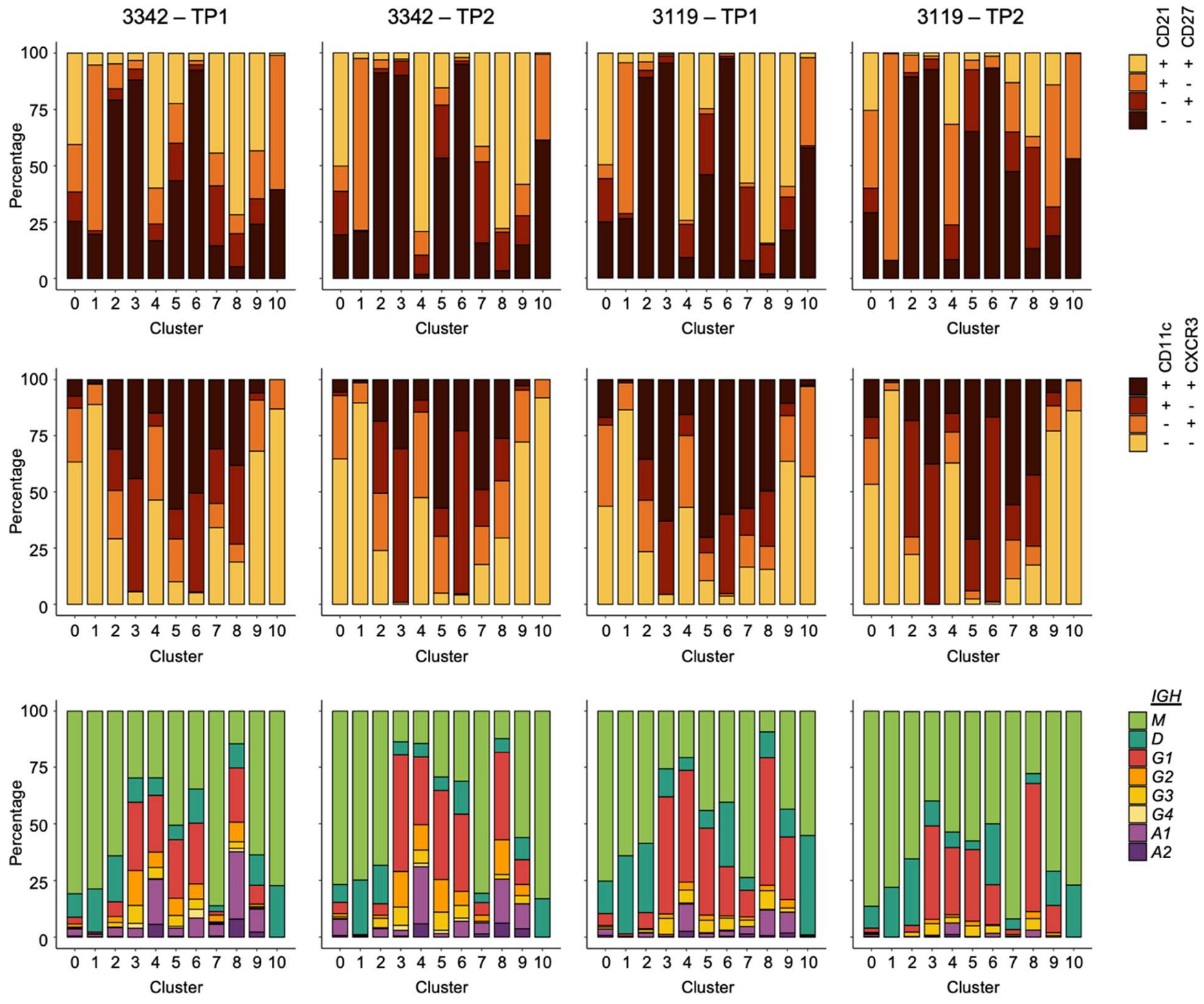


**Figure S5: Distribution of cell surface marker expression and immunoglobulin heavy chain transcripts per transcriptional cluster in each sample.** The top row shows the percentage of CD21$^{-/+}$ CD27$^{-/+}$ B cells, the middle row shows the percentage of CD11c$^{-/+}$ CXCR3$^{-/+}$ B cells, and the bottom row shows the percentage of immunoglobulin heavy chain transcripts.

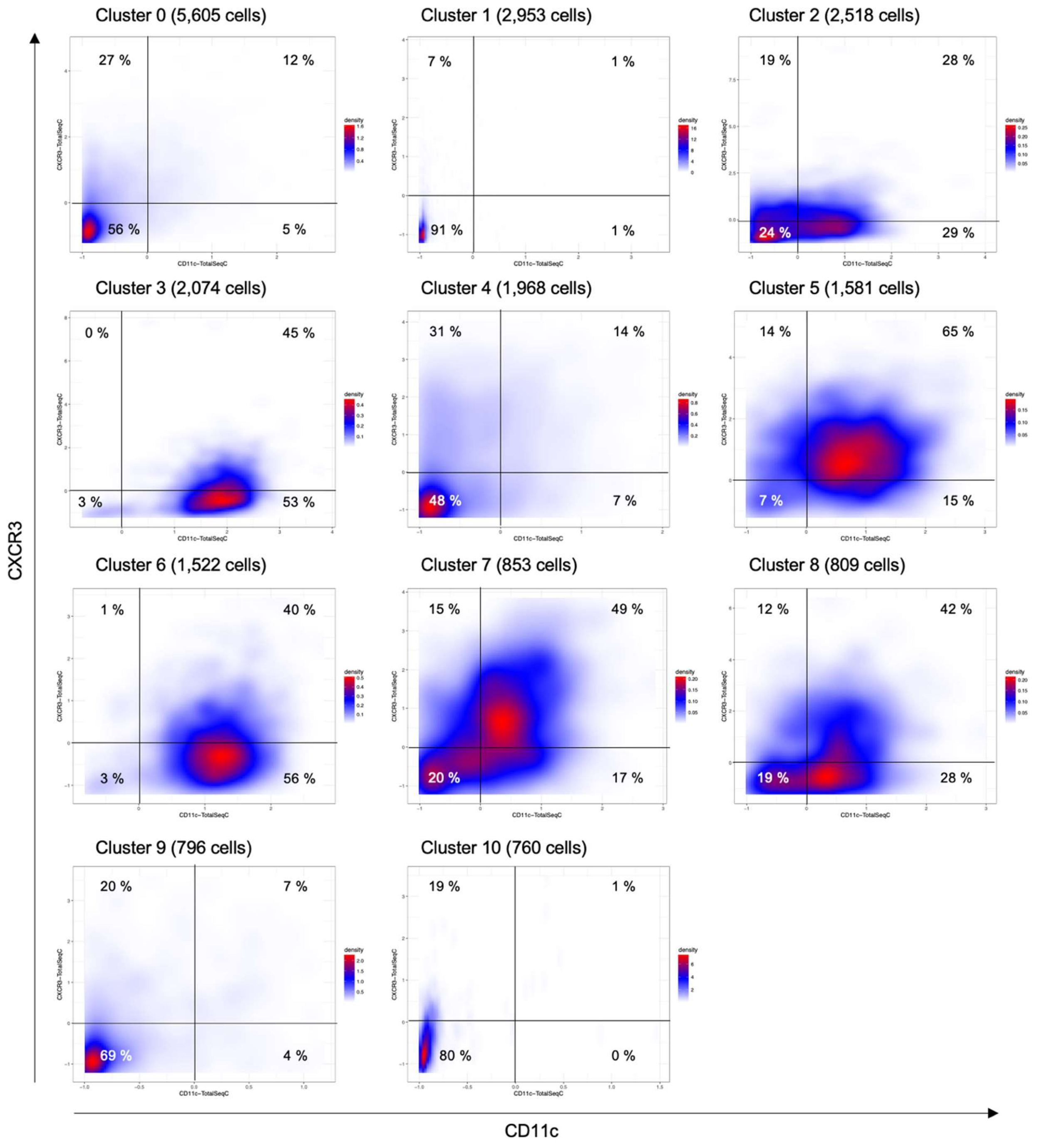


**Figure S6: Cell surface expression of CD11c and CXCR3 in each cluster as determined by feature barcoding.** Lines in the graphs denote the cutoffs for expression (normalized values > 0).

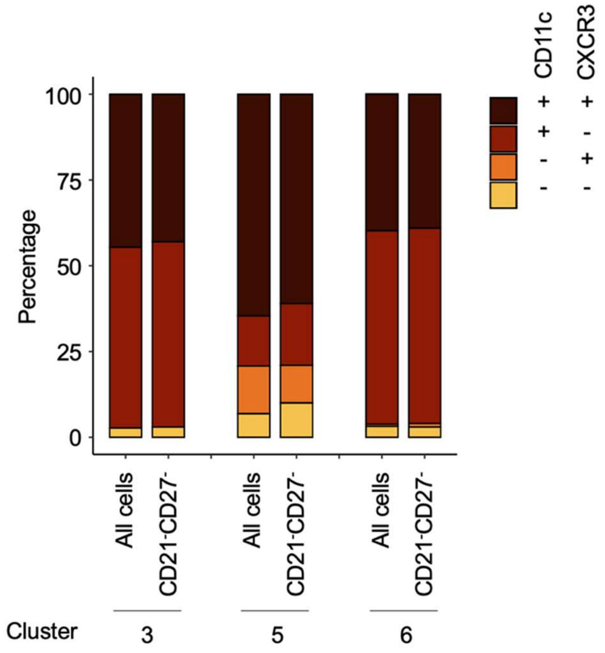


**Figure S7: Distribution of CD11c and CXCR3 among all cells and CD21$^-$CD27$^-$ cells in the three clusters that contain atypical B cells.**

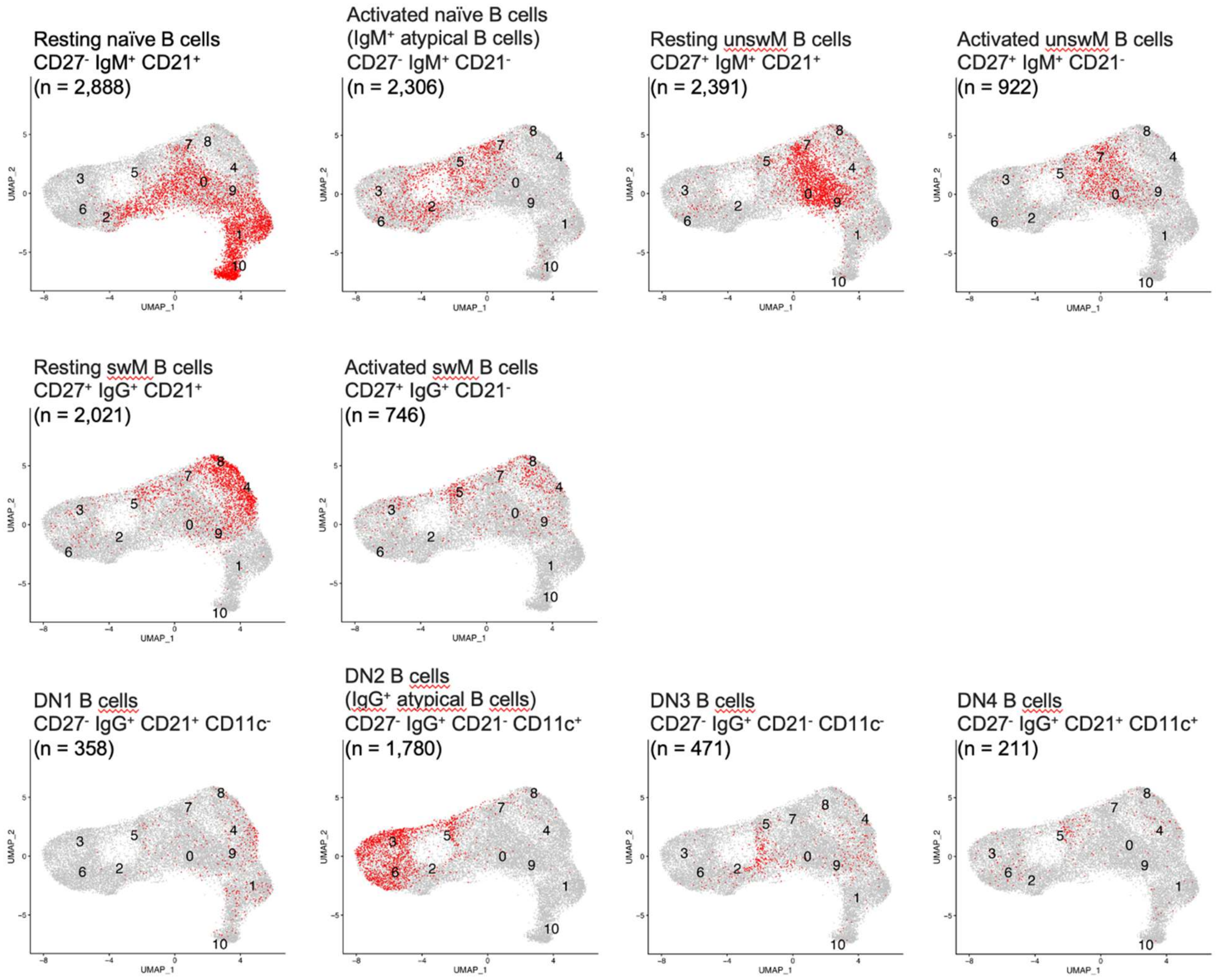


**Figure S8: Phenotypically defined B cell subsets projected onto the transcriptomics-based UMAP.** B cell subsets were defined according to published guidelines (*22*), with modifications. IgD, IgA, and IgE were not included in our panel of surface markers. We therefore used IgM to identify naïve and unswitched memory (unswM) B cells and IgG to define class-switched memory (swM) and double negative (DN) B cells. Cells of the indicated phenotype are shown in red, while all other B cells are shown in gray.

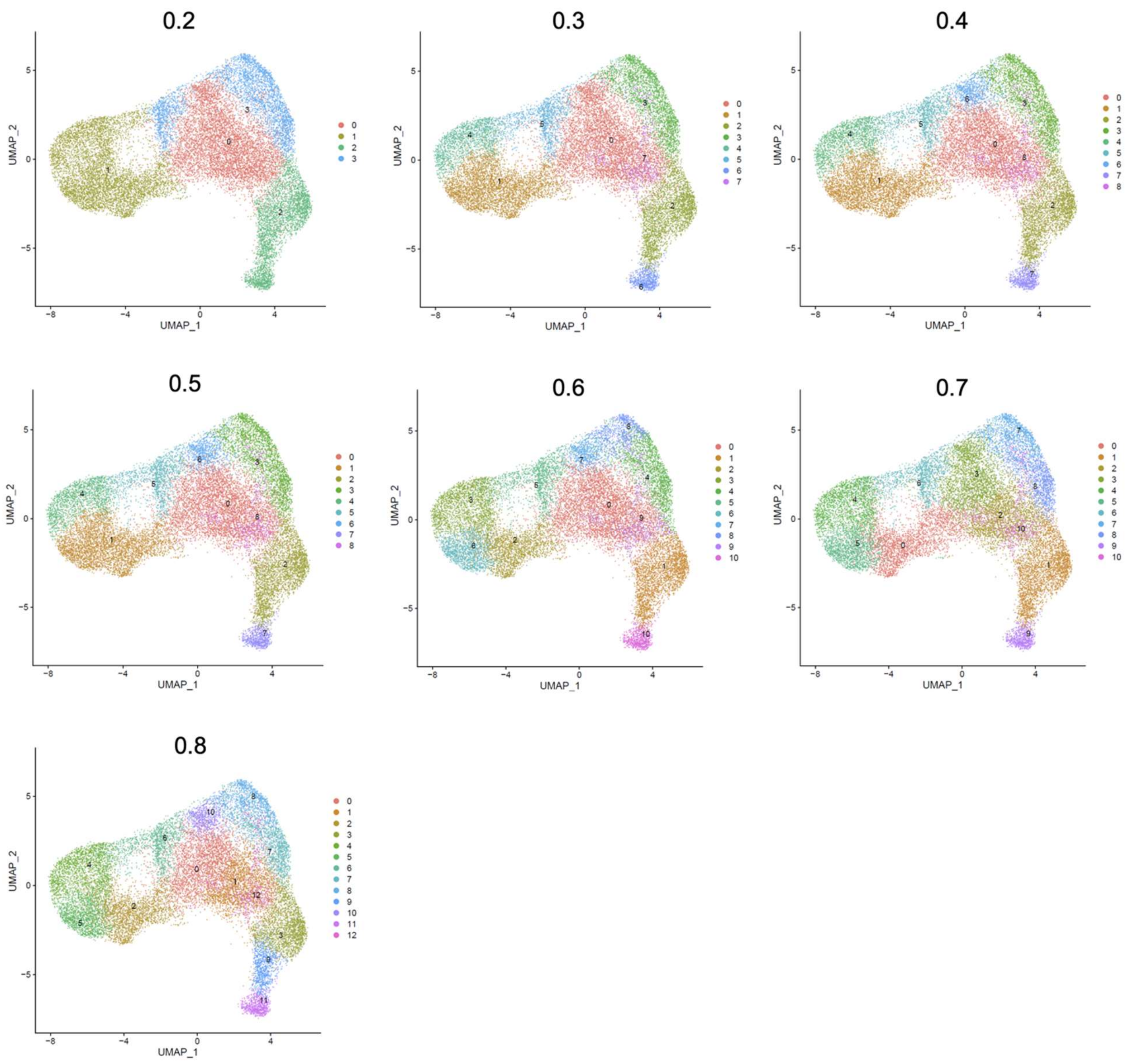


**Figure S9: Clusters obtained at a range of resolutions in Seurat.** The results of clustering at a resolution of 0.5 were used in this study.

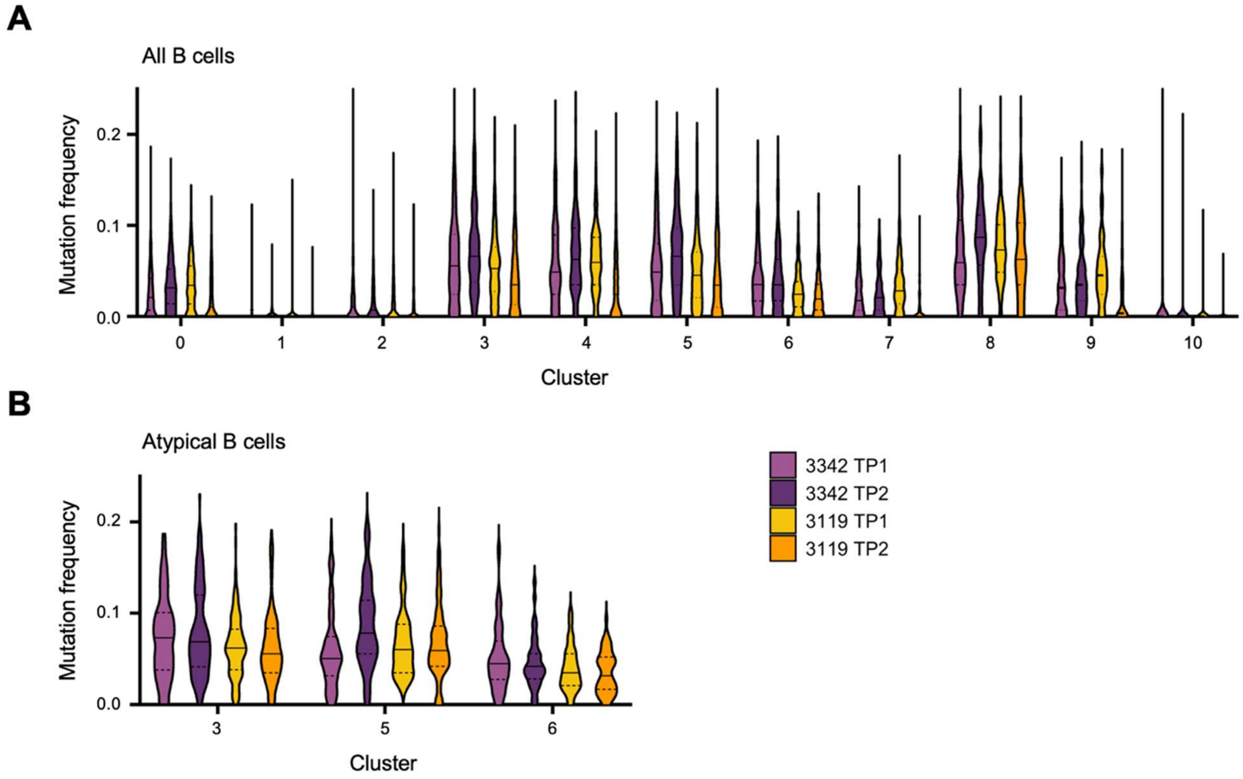


**Figure S10: Levels of somatic hypermutation in each cluster per sample. A)** Levels of somatic hypermutation in all clusters. **B)** Levels of somatic hypermutation in atypical B cells of clusters 3, 5, and 6.

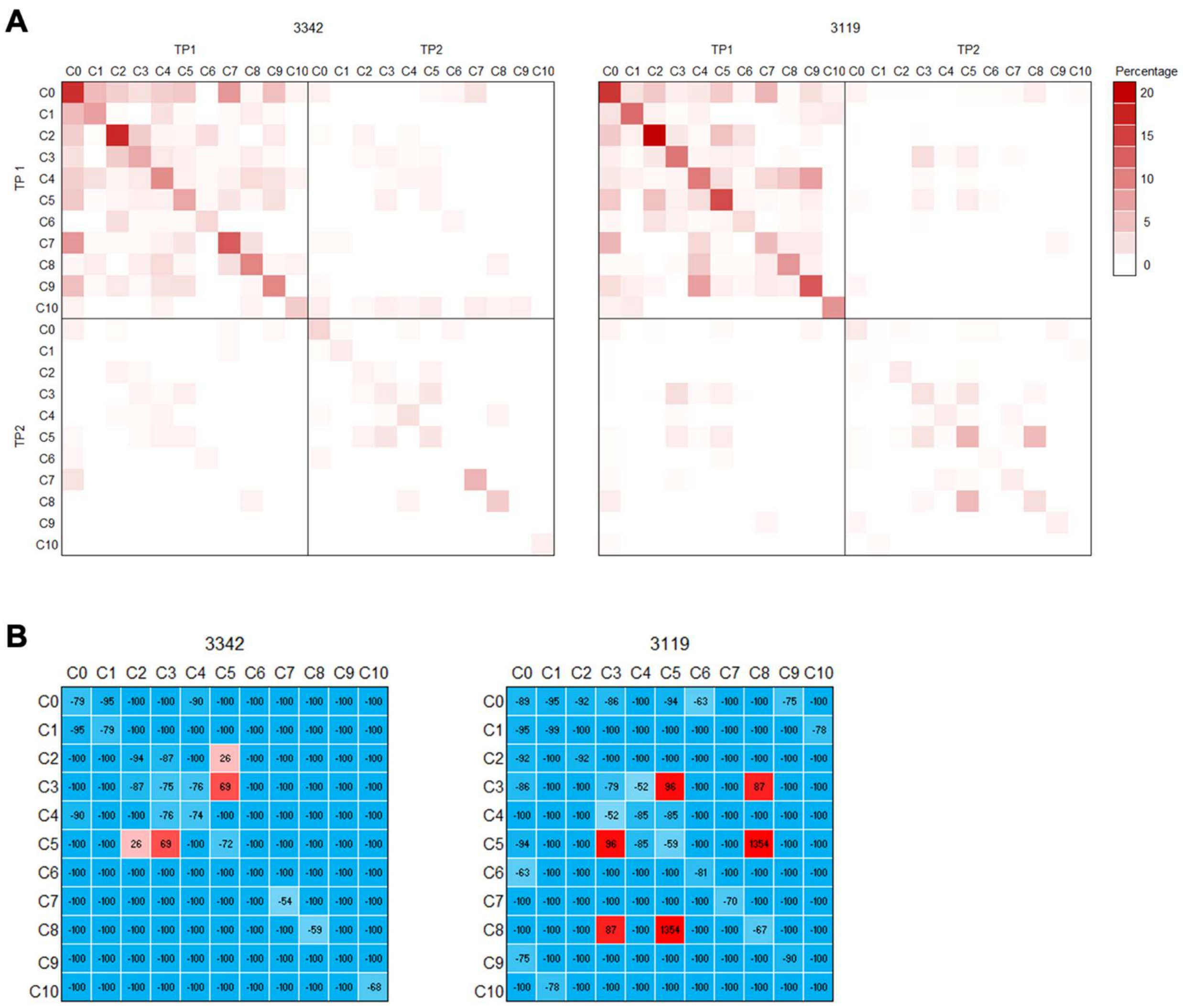


3342

| | C0 | C1 | C2 | C3 | C4 | C5 | C6 | C7 | C8 | C9 | C10 |
|---|---|---|---|---|---|---|---|---|---|---|---|
| C0 | -79 | -95 | -100 | -100 | -90 | -100 | -100 | -100 | -100 | -100 | -100 |
| C1 | -95 | -79 | -100 | -100 | -100 | -100 | -100 | -100 | -100 | -100 | -100 |
| C2 | -100 | -100 | -94 | -87 | -100 | 26 | -100 | -100 | -100 | -100 | -100 |
| C3 | -100 | -100 | -87 | -75 | -76 | 69 | -100 | -100 | -100 | -100 | -100 |
| C4 | -90 | -100 | -100 | -76 | -74 | -100 | -100 | -100 | -100 | -100 | -100 |
| C5 | -100 | -100 | 26 | 69 | -100 | -72 | -100 | -100 | -100 | -100 | -100 |
| C6 | -100 | -100 | -100 | -100 | -100 | -100 | -100 | -100 | -100 | -100 | -100 |
| C7 | -100 | -100 | -100 | -100 | -100 | -100 | -100 | -54 | -100 | -100 | -100 |
| C8 | -100 | -100 | -100 | -100 | -100 | -100 | -100 | -100 | -59 | -100 | -100 |
| C9 | -100 | -100 | -100 | -100 | -100 | -100 | -100 | -100 | -100 | -100 | -100 |
| C10 | -100 | -100 | -100 | -100 | -100 | -100 | -100 | -100 | -100 | -100 | -68 |

3119

| | C0 | C1 | C2 | C3 | C4 | C5 | C6 | C7 | C8 | C9 | C10 |
|---|---|---|---|---|---|---|---|---|---|---|---|
| C0 | -89 | -95 | -92 | -86 | -100 | -94 | -63 | -100 | -100 | -75 | -100 |
| C1 | -95 | -99 | -100 | -100 | -100 | -100 | -100 | -100 | -100 | -100 | -78 |
| C2 | -92 | -100 | -92 | -100 | -100 | -100 | -100 | -100 | -100 | -100 | -100 |
| C3 | -86 | -100 | -100 | -79 | -52 | 96 | -100 | -100 | 87 | -100 | -100 |
| C4 | -100 | -100 | -100 | -52 | -85 | -85 | -100 | -100 | -100 | -100 | -100 |
| C5 | -94 | -100 | -100 | 96 | -85 | -59 | -100 | -100 | 1354 | -100 | -100 |
| C6 | -63 | -100 | -100 | -100 | -100 | -100 | -81 | -100 | -100 | -100 | -100 |
| C7 | -100 | -100 | -100 | -100 | -100 | -100 | -100 | -70 | -100 | -100 | -100 |
| C8 | -100 | -100 | -100 | 87 | -100 | 1354 | -100 | -100 | -67 | -100 | -100 |
| C9 | -75 | -100 | -100 | -100 | -100 | -100 | -100 | -100 | -100 | -90 | -100 |
| C10 | -100 | -78 | -100 | -100 | -100 | -100 | -100 | -100 | -100 | -100 | -100 |

**Figure S11: Clonal expansion within and clonal connections between all transcriptomics-based clusters. A)** Clonal expansion within clusters (along the diagonal in each heatmap) and clonal connections between clusters (off-diagonal cells). **B)** The percent difference in clonal expansion/connections between the first and the second time point.

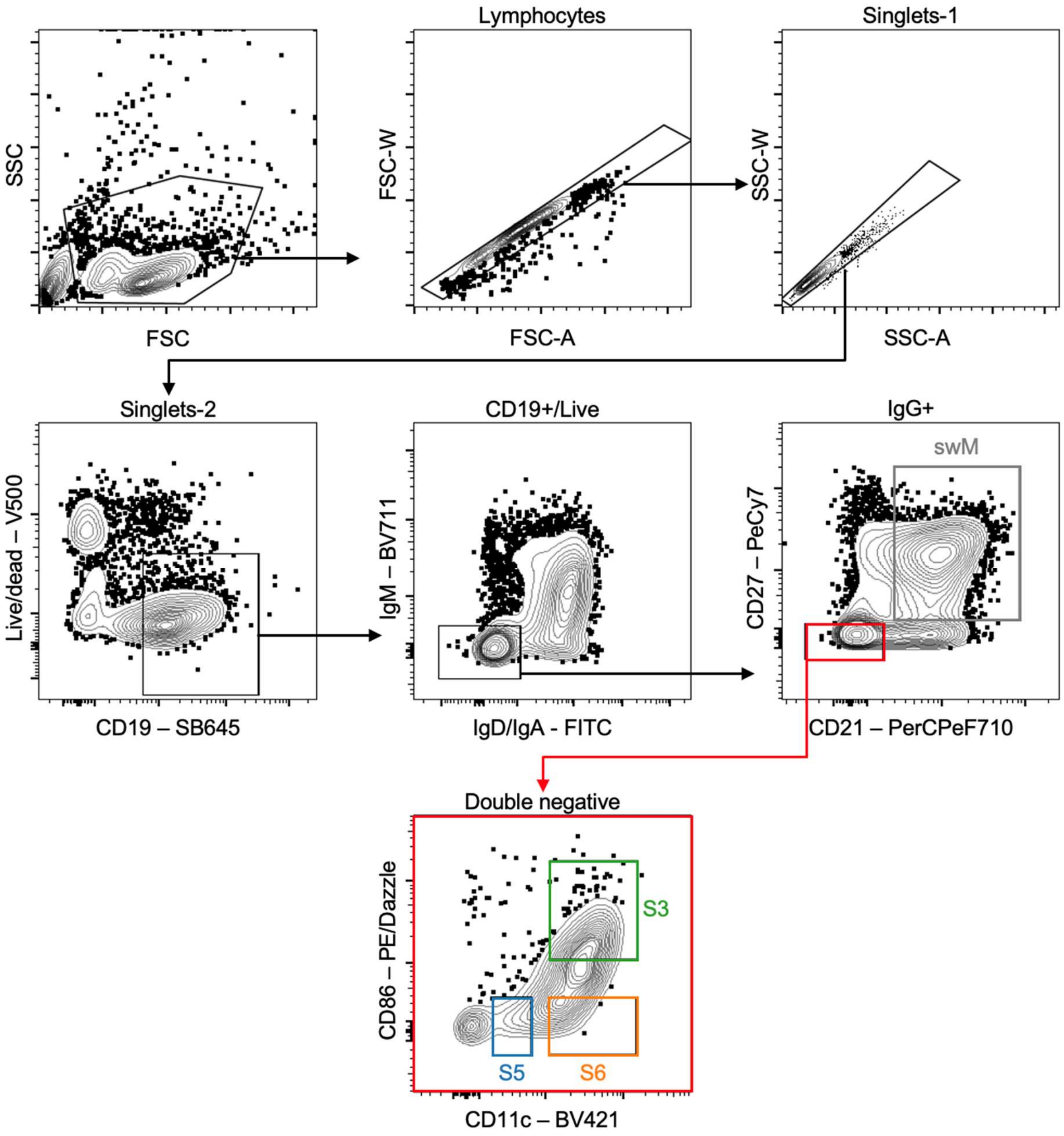


**Figure S12: Gating strategy used to sort atypical B cell subsets.**

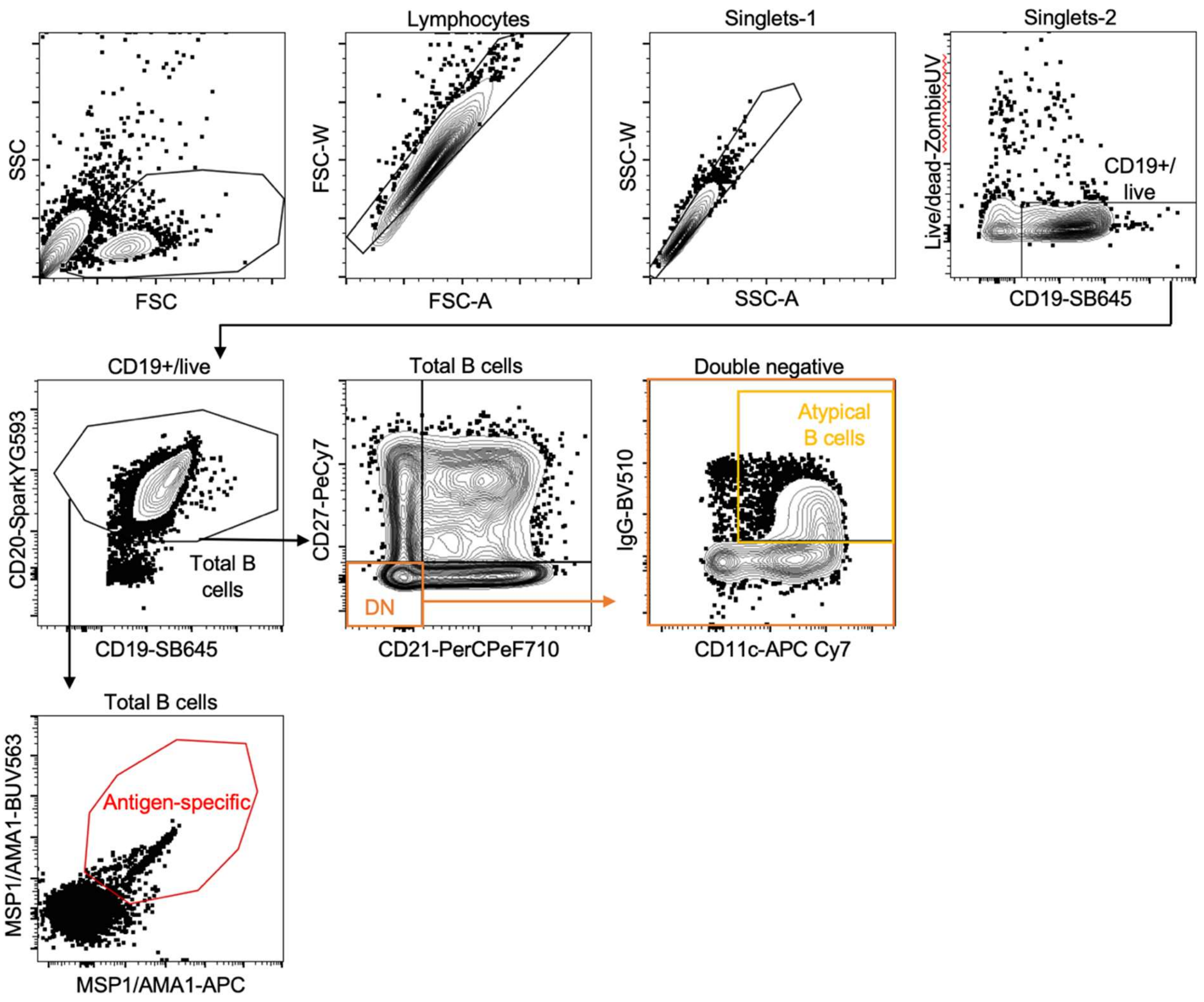


**Figure S13: Gating strategy used to identify atypical B cells and antigen-specific B cells for spectral flow analysis.**

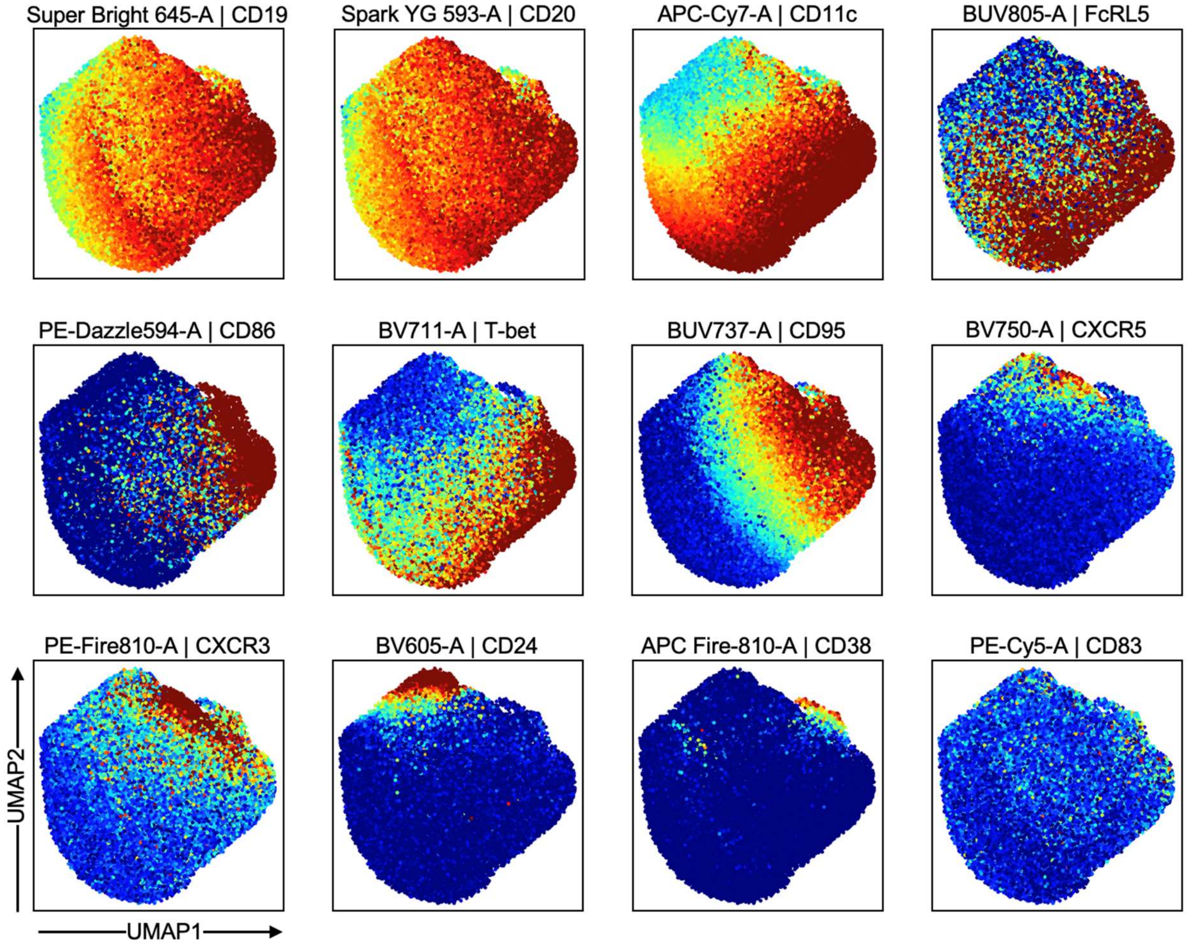


**Figure S14: Projection of the 12 surface and intracellular markers used to generate the composite UMAP of atypical B cells onto this UMAP.**

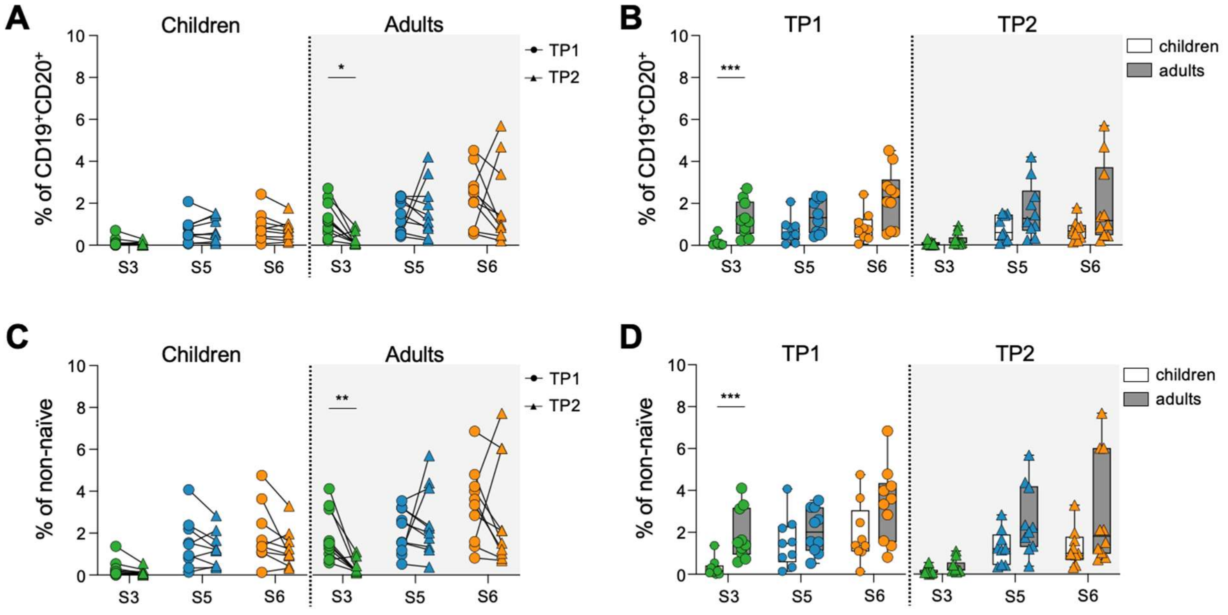


**Figure S15: Percentage of atypical B cell subsets among total and non-naïve B cells. A)** Percentage of each atypical B cell subset in children (n = 9) and adults (n = 10) at the first time point (TP1; circles) and second time point (TP2; triangles) among total $CD19^{+}CD20^{+}$ B cells. Data points from the same individual are connected with a black line. **B)** Comparison of the percentage of atypical B cell subsets among $CD19^{+}CD20^{+}$ B cells between children and adults at each time point. **C)** Percentage of each atypical B cell subset in children and adults at the first time point (TP1; circles) and second time point (TP2; triangles) among non-naïve B cells (total B cells with the exception of naïve $IgD^{+}CD27^{-}$ B cells). Data points from the same individual are connected with a black line. **D)** Comparison of the percentage of atypical B cell subsets among non-naïve B cells between children and adults at each time point. Statistical analysis was performed using a Wilcoxon matched-pairs signed rank test for paired data or a Mann-Whitney test for unpaired data with a 10% false discovery rate using the two-stage step-up method of Benjamini, Krieger, and Yekutieli. *P < 0.05, **P < 0.01, ***P < 0.001

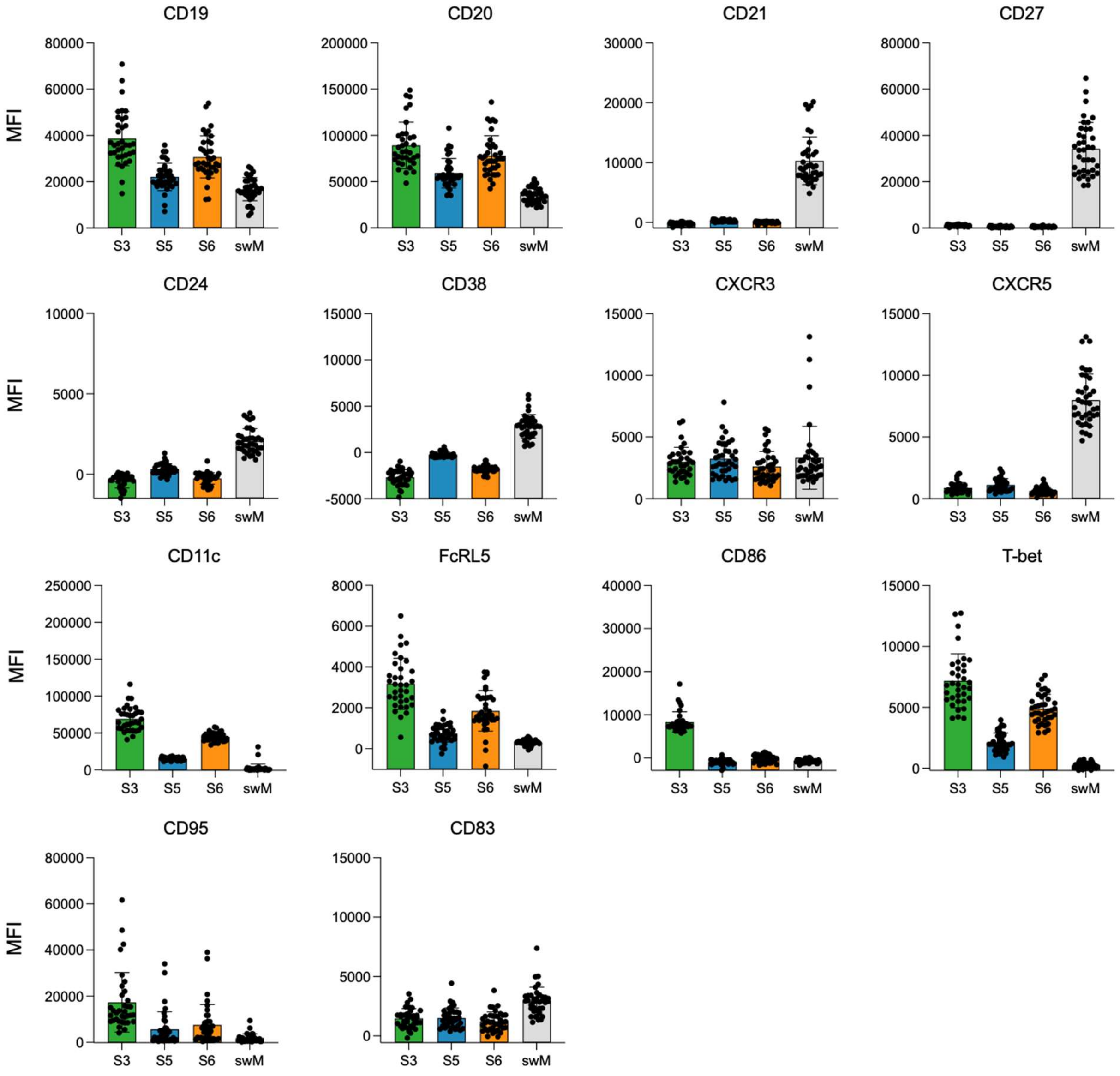


**Figure S16: Mean fluorescence intensity (MFI) of cell surface and intracellular markers used for the flow cytometric analysis of atypical B cell subsets.** S3, subset 3; S5, subset 5; S6, subset 6; swM, resting switched memory B cells.

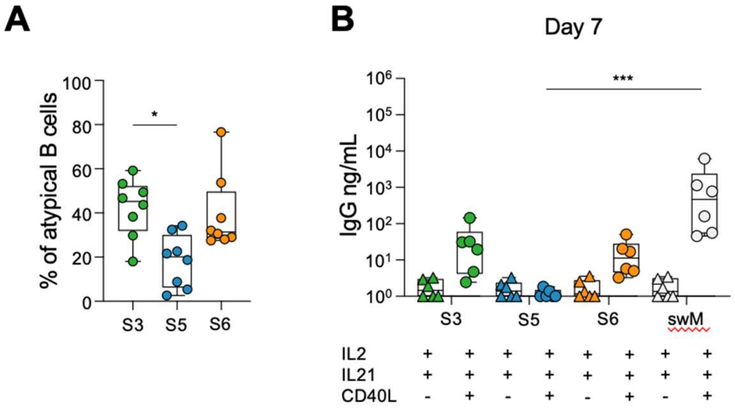


**Figure S17: IgG production among three atypical B cell subsets. A)** Distribution of atypical B cells over the three subsets. **B)** IgG concentration in culture supernatant after 14 days of in vitro culture. A one-way ANOVA was used to test for statistically significant differences between the groups. P values shown are from Kruskal-Wallis post hoc test. ** $P < 0.01$; *** $P < 0.001$; **** $P < 0.0001$.

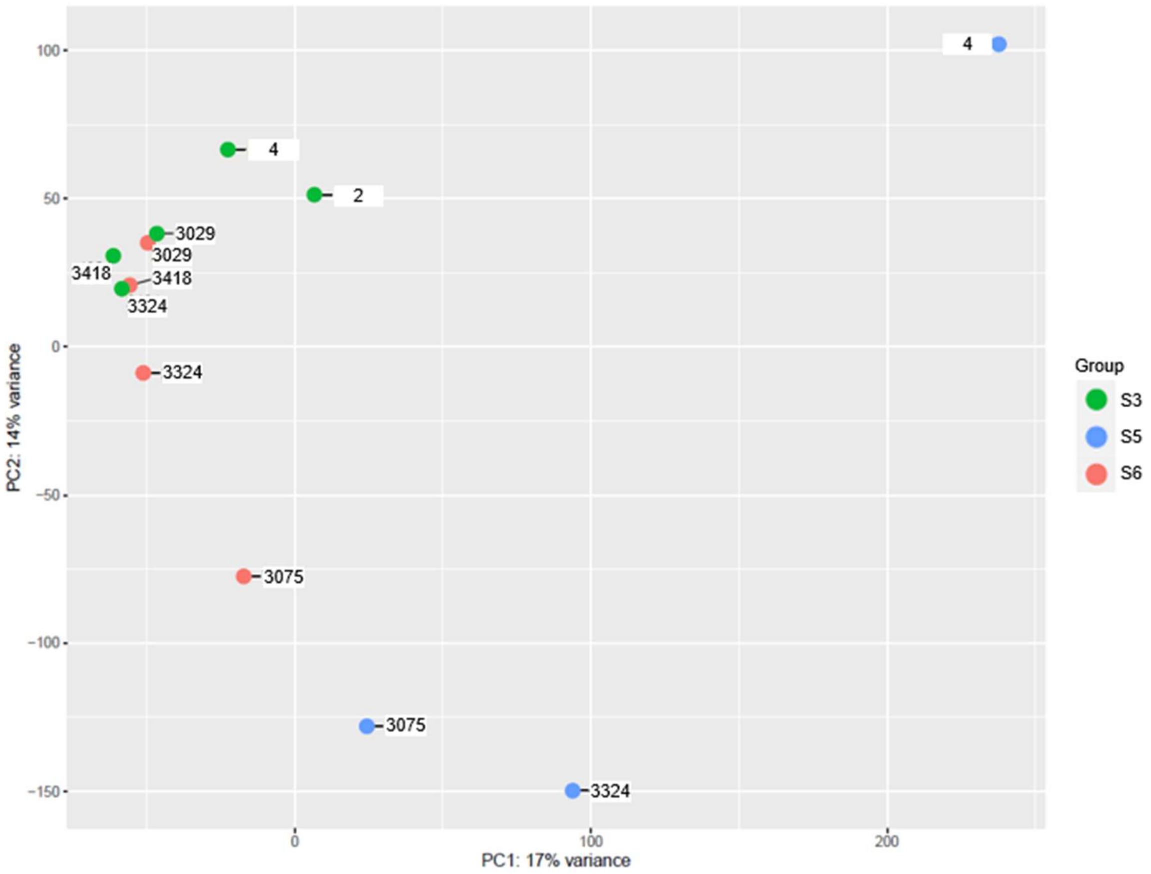


**Figure S18: Principal component analysis plot of the 5000 most variable genes among bulk atypical B cell subsets.** For each subset, RNA-seq libraries were prepared from 200 cells that were sorted by flow cytometry based on the expression of CD11c and CD86. Each sample is labeled with the donor ID and color-coded by subset.

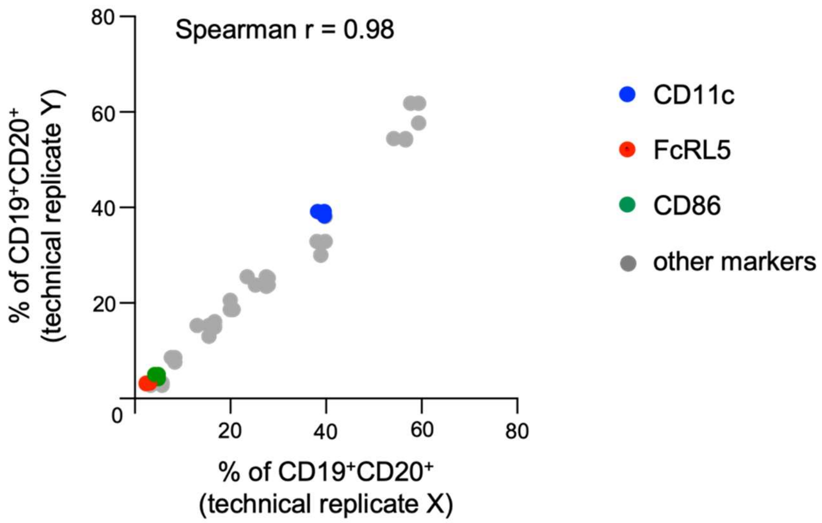


**Figure S19: Spectral flow cytometry technical replicates.** The frequency of cells positive for the markers CD11c, FcRL5, CD86 and all other surface and intracellular markers used in the study, with the exception of CD19 and CD20, are shown.

**Table S1: Information about all individuals included in the study**. Provided as separate Excel file.

**Table S2: Number of cells sorted and pooled for quality control**

| Donor | Time point | Number of sorted cells | | Number of pooled cells | |
|---|---|---|---|---|---|
| | | Antigen-experienced B cells[1] | Naïve B cells[2] | Antigen-experienced B cells | Naïve B cells |
| 3342 | 1 | 28,519 | 24,289 | 28,519 | 21,481 |
| 3342 | 2 | 42,062 | 48,133 | 42,062 | 7,938 |
| 3119 | 1 | 62,035 | 94,536 | 45,000 | 5,000 |
| 3119 | 2 | 32,098 | 23,851 | 32,098 | 17,902 |
| Average of total B cells (%) | | 49 | 51 | 74 | 26 |

[1] Antigen-experienced B cells were defined as $CD21^-CD27^-$ atypical B cells, $CD21^-CD27^+$ activated B cells, and $CD21^+CD27^+$ memory B cells

[2] Naïve B cells were defined as $CD21^+CD27^-$

**Table S3: Summary of sequence stats**

| Donor | Time point | Sequence saturation (%) | Output from Cell Ranger | | | # cells after filtering |
|---|---|---|---|---|---|---|
| | | | # cells before filtering | Mean reads per cell | Median genes per cell | |
| 3342 | 1 | 89.6 | 8,899 | 63,791 | 893 | 6,070 |
| 3342 | 2 | 90.1 | 6,773 | 82,555 | 1,160 | 3,399 |
| 3119 | 1 | 93.3 | 6,726 | 87,583 | 982 | 6,067 |
| 3119 | 2 | 92.8 | 7,018 | 97,320 | 1,167 | 5,903 |
| Total number of cells that passed quality control: | | | | | | 21,439 |

**Table S4: Numbers of cells per sample in each cluster**

| Donor | Time point | Cluster | | | | | | | | | | |
|---|---|---|---|---|---|---|---|---|---|---|---|---|
| | | 0 | 1 | 2 | 3 | 4 | 5 | 6 | 7 | 8 | 9 | 10 |
| 3342 | 1 | 2058 | 731 | 526 | 548 | 642 | 448 | 331 | 214 | 191 | 282 | 99 |
| 3342 | 2 | 957 | 404 | 259 | 361 | 399 | 285 | 197 | 147 | 122 | 108 | 160 |
| 3119 | 1 | 1499 | 518 | 1032 | 529 | 615 | 370 | 529 | 218 | 329 | 236 | 192 |
| 3119 | 2 | 1091 | 1300 | 701 | 636 | 312 | 478 | 465 | 274 | 167 | 170 | 309 |

**Table S5: Differentially expressed genes between clusters 3, 5, and 6**. Provided as separate Excel file.

**Table S6: Reactome pathway analysis results**. Provided as separate Excel file.

**Table S7: Antibodies used for flow sorting and single cell sequencing**

| Antibody | Clone | Company | Catalog number |
|---|---|---|---|
| Super Bright 600 anti-human CD10 | CB-CALLA | Thermo | 63-0106-41 |
| Brilliant Violet 421 anti-human CD19 | SJ25C1 | BioLegend | 363017 |
| Brilliant Violet 785 anti-human CD20 | 2H7 | BioLegend | 302355 |
| PerCP-eFluor 710 anti-human CD21 | HB5 | Thermo | 46-0219-41 |
| PE/Cyanine7 anti-human CD27 | O323 | BioLegend | 302837 |
| TotalSeq-C0181 anti-human CD21 | Bu32 | BioLegend | 354923 |
| TotalSeq-C0154 anti-human CD27 | O323 | BioLegend | 302853 |
| TotalSeq-C0053 anti-human CD11c | S-HCL-3 | BioLegend | 371521 |
| TotalSeq-C0140 anti-human CXCR3 | G025H7 | BioLegend | 353747 |
| TotalSeq-C0375 anti-human IgG | M1310G05 | BioLegend | 410727 |
| TotalSeq-C0136 anti-human IgM | MHM-88 | BioLegend | 314547 |

**Table S8:** Reagents used for single-cell immune profiling and Feature Barcoding

| Reagent | Company | Catalog number |
|---|---|---|
| Human TruStain FcX | BioLegend | 422301 |
| Dextran sulfate sodium salt | Sigma-Aldrich | 42867-5G |
| Chromium Single Cell 5' Library & Gel Bead Kit v1 | 10x Genomics | 1000006 |
| Chromium Single Cell A Chip Kit | 10x Genomics | 1000151 |
| Chromium Single Cell 5' Feature Barcode Library Kit | 10x Genomics | 1000080 |
| Chromium Single Cell 5' Library Construction Kit | 10x Genomics | 1000020 |
| Chromium Single Cell V(D)J Enrichment Kit | 10x Genomics | 1000016 |
| Agencourt AMPure XP beads | Beckman Coulter | A63880 |

**Table S9: Average gene expression values for all genes in all clusters**. Provided as a separate Excel file.

**Table S10: Streptavidin conjugates used for spectral flow cytometry**

| Antigen | Streptavidin-fluor | Company | Catalog number |
|---|---|---|---|
| Biotinylated merozoite proteins | APC | Tonbo | 20-4317-U100 |
| Biotinylated merozoite proteins | BUV563 | BD | 612935 |

**Table S11: Antibodies used in spectral flow cytometry analysis**

| Antibody | Clone | Company | Catalog number |
|---|---|---|---|
| Super Bright 645 anti-human CD19 | HIB19 | Invitrogen | 64019942 |
| Spark YG 593 anti-human CD20 | 2H7 | BioLegend | 302367 |
| PerCP-eFluor 710 anti-human CD21 | HB5 | Thermo | 46-0219-42 |
| Brilliant Violet 605 anti-human CD24 | ML5 | BioLegend | 311123 |
| PE/Cyanine7 anti-human CD27 | O323 | BioLegend | 302837 |
| APC/Fire 810 anti-human CD38 | HB-7 | BioLegend | 356643 |
| PE/Cyanine5 anti-human CD83 | HB15e | BioLegend | 305310 |
| PE-Dazzle 594 anti-human CD86 | BU63 | BioLegend | 374217 |
| BUV737 anti-human CD95 | DX2 | BD | 612790 |
| Pacific Blue anti-human IgD | IA6-2 | BioLegend | 348223 |
| BV570 anti-human IgM | MHM-88 | BioLegend | 314517 |
| BV510 anti-human IgG | M1310G05 | BioLegend | 410715 |
| FITC anti-human IgA | mA-6E1 | Miltenyi | 130114001 |
| PE-Fire 810 anti-human CXCR3 | G025H7 | BioLegend | 353759 |
| Brilliant Violet 750 anti-human CXCR5 | J252D4 | BioLegend | 356941 |
| APC/Cyanine 7 anti-human CD11c | Bu15 | BioLegend | 337217 |
| Brilliant Violet 711 anti-human Tbet (used for intracellular staining step) | 4B10 | BioLegend | 644819 |
| BUV805 anti-human FcRL5 | 509F6 | BD | 749599 |
